# A theoretical model for focal seizure initiation, propagation, termination, and progression

**DOI:** 10.1101/724088

**Authors:** Jyun-you Liou, Elliot H. Smith, Lisa M. Bateman, Samuel L. Bruce, Guy M. McKhann, Robert R. Goodman, Ronald G. Emerson, Catherine A. Schevon, L. F. Abbott

**Affiliations:** Department of Physiology and Cellular Biophysics, Columbia University, New York, NY 10032, USA; Department of Anesthesiology, NewYork-Presbyterian Hospital/Weill Cornell Medicine, New York, NY 10065, USA; Department of Neurology, Columbia University Medical Center, New York, New York 10032, USA; Department of Neurological Surgery, Columbia University Medical Center, New York, NY, 10032, USA; Vagelos College of Physicians & Surgeons, Columbia University, New York, NY 10032; Mortimer B. Zuckerman Mind Brain Behavior Institute, Department of Neuroscience, Columbia University, New York, NY 10027, USA; Hospital for Special Surgery, Weill Cornell Medical College, 535 East 70th Street, New York, NY; Department of Neurosurgery, Icahn School of Medicine at Mount Sinai, 1 West 85th St., Suite 1C, New York, NY; Department of Neurosurgery, University of Utah, 175 N Medical Dr E #5, Salt Lake City, UT

## Abstract

We developed a neural network model that can account for the major elements common to human focal seizures. These include the tonic-clonic transition, slow advance of clinical semiology and corresponding seizure territory expansion, widespread EEG synchronization, and slowing of the ictal rhythm as the seizure approaches termination. These were reproduced by incorporating usage-dependent exhaustion of inhibition in an adaptive neural network that receives global feedback inhibition in addition to local recurrent projections. Our model proposes mechanisms that may underline common EEG seizure onset patterns and status epilepticus and postulates a role for synaptic plasticity in emergence of epileptic foci. Complex patterns of seizure activity and bi-stable seizure evolution end-points arise when stochastic noise is included. With the rapid advancement of clinical and experimental tools, we believe that this can provide a roadmap and potentially a testbed for future explorations of seizure mechanisms and clinical therapies.

## Introduction

Focal seizures have been recognized for more than 3000 years, with descriptions dating back to ancient Mesopotamia (Worthington, 2007). Although focal seizures can present with a plethora of behavioral manifestations that vary according to the affected cortical regions, there are several consistent clinical and large-scale EEG features (Kotagal and Luders, 2008): propagation from a focal onset location to large brain regions, widespread neuronal synchronization, a transition from tonic to clonic activity, and a slowing pace of neuronal discharging prior to simultaneous seizure termination.

Our recent investigations, utilizing microelectrode array recordings in humans, identified multiunit and single unit underpinnings of these common seizure features (Schevon *et al*., 2012; Smith *et al*., 2016). Based on these findings and results of parallel animal model studies (Trevelyan *et al*., 2006; Trevelyan *et al*., 2007a; Trevelyan *et al*., 2007b), we proposed a dual spatial structure for focal seizures consisting of a core region of seizing brain bounded by an ictal wavefront surrounded by a passively reactive penumbra. What delineates the boundary of the ictal core is the ictal wavefront, a narrow band of intense, desynchronized multiunit (tonic) firing that marks the transition to seizure at a given brain location. Typically, this tonic firing phase is not detectable in clinical EEG or band-limited local field potentials (LFPs). Evidence from human (Schevon *et al*., 2012) and animal studies (Trevelyan *et al*., 2006; Trevelyan *et al*., 2007b) suggests that collapse of inhibition is the key element causing ictal wavefront propagation, which leads to progressive seizure territory expansion. The slow pace of ictal wavefront propagation (< 1 mm/sec) corresponds to the slow evolution of the electrographic seizure and clinical semiology, e.g. the classic Jacksonian march (York and Steinberg, 2011). As it advances, the ictal wavefront generates fast-moving ictal discharges (Trevelyan, 2009; Smith *et al*., 2016), directed inward towards the initiation point, at speeds two orders of magnitude higher than the wavefront propagation (Smith *et al*., 2016; Liou *et al*., 2017). These fast-moving ictal discharges are the basis of the well-recognized high amplitude field potential deflections that are the hallmark of electrographic seizures. Thus, there are two types of moving waves that characterize seizures: fast, inward-moving ictal discharges and the slow outward-moving wave of ictal recruitment.

To date, despite extensive prior work in computational seizure modeling (Soltesz and Staley, 2011), no theoretical study has shown how this dynamic topological structure can arise from basic epileptic mechanisms. Phenomenological models, such as Epileptor (Jirsa *et al*., 2014; Proix *et al*., 2018), have successfully utilized large-scale functional EEG features to reproduce the large-area, apparently synchronized EEG activity that characterizes clinical seizure recordings, although identifying the neurophysiological processes corresponding to each abstract variable can be a challenging task. Biophysical models have been used to explore the role of specific neurophysiological processes responsible for seizure transitions, such as intracellular and extracellular potassium dynamics (Cressman *et al*., 2009; Ullah *et al*., 2009), transmembrane chloride gradients (Buchin *et al*., 2016), and calcium-activated processes (Yang *et al*., 2005). However, the evolving, wide-area dynamics that characterize the life cycle of a seizure (Smith *et al*., 2016) remain difficult to explain by any single biophysical mechanism. Moreover, given that seizures can simultaneously activate a multitude of pathophysiological processes across extensive interconnected brain regions, biophysical models may be limited to a fragmented or overly specific account of seizure dynamics.

Here, we describe a biophysically-constrained cortical network model designed to link the key pathological cellular mechanisms that underpin seizures to their large-scale spatial structure. Inspired by the spirit of phenomenological models, we adopt an approach with minimal assumptions, aiming to show that complex spatiotemporal dynamics can arise from simple, generalizable, and experimentally validated biophysical principles. We demonstrate that maintaining the normal transmembrane chloride gradient is critical for inhibition robustness, which is necessary for restricting seizure propagation. Our theory provides a consistent framework explaining the key clinical features widely observed in focal seizure patients (Kotagal and Luders, 2008; Ebersole *et al*., 2014; Extercatte *et al*., 2015). We also test several predictions arising from the model using our existing dataset of microelectrode recordings of seizures in epilepsy patients.

## Results

We modeled the neocortex as a 2-dimensional neuron sheet (Bressloff, 2014). Model neurons are pyramidal cell-like and contain two intrinsic conductances – leak and slow potassium – and two synaptic conductances – excitatory (AMPA) and inhibitory (GABA-A). Neuronal mean firing rates are calculated by passing the difference between membrane potentials and thresholds through a sigmoid function. Model neurons incorporate spike-frequency adaptation mechanisms – neuronal firing increases the spike threshold and activates an additional slow afterhyperpolarization (sAHP) conductance. Neurons are recurrently connected by both excitatory and inhibitory synaptic projections (Figure 1A) (Prince and Wilder, 1967; Kramer *et al*., 2005). Intra-cortical projections are distance-dependent, with the range of inhibitory projections longer than the excitatory range, thereby making the spatial distribution of synaptic weights follow a “Mexican hat” structure (Prince and Wilder, 1967; Coombes, 2005; Bressloff, 2014). In addition, our model includes a distance-independent recurrent inhibition pathway (Figure 1A, part γ), inspired by recent observation of large-scale inhibitory effects of focal seizure activity (Liou *et al*., 2018). Interneuron membrane potential dynamics are not directly modelled. Instead, interneurons are implicitly incorporated through the recurrent inhibitory projections (Supplementary Online Methods, Equation 5). Biophysical parameters are set based on previously reported values from recorded neocortical pyramidal neurons (Supplementary Online Methods, Table S1). In its original form, the model does not spontaneously seize, instead model seizures are initiated by transient focal excitatory input. Thus it can be considered a model of a health neural circuit, with the degree of resistance to seizure controlled by model parameters.

**Figure 1.**
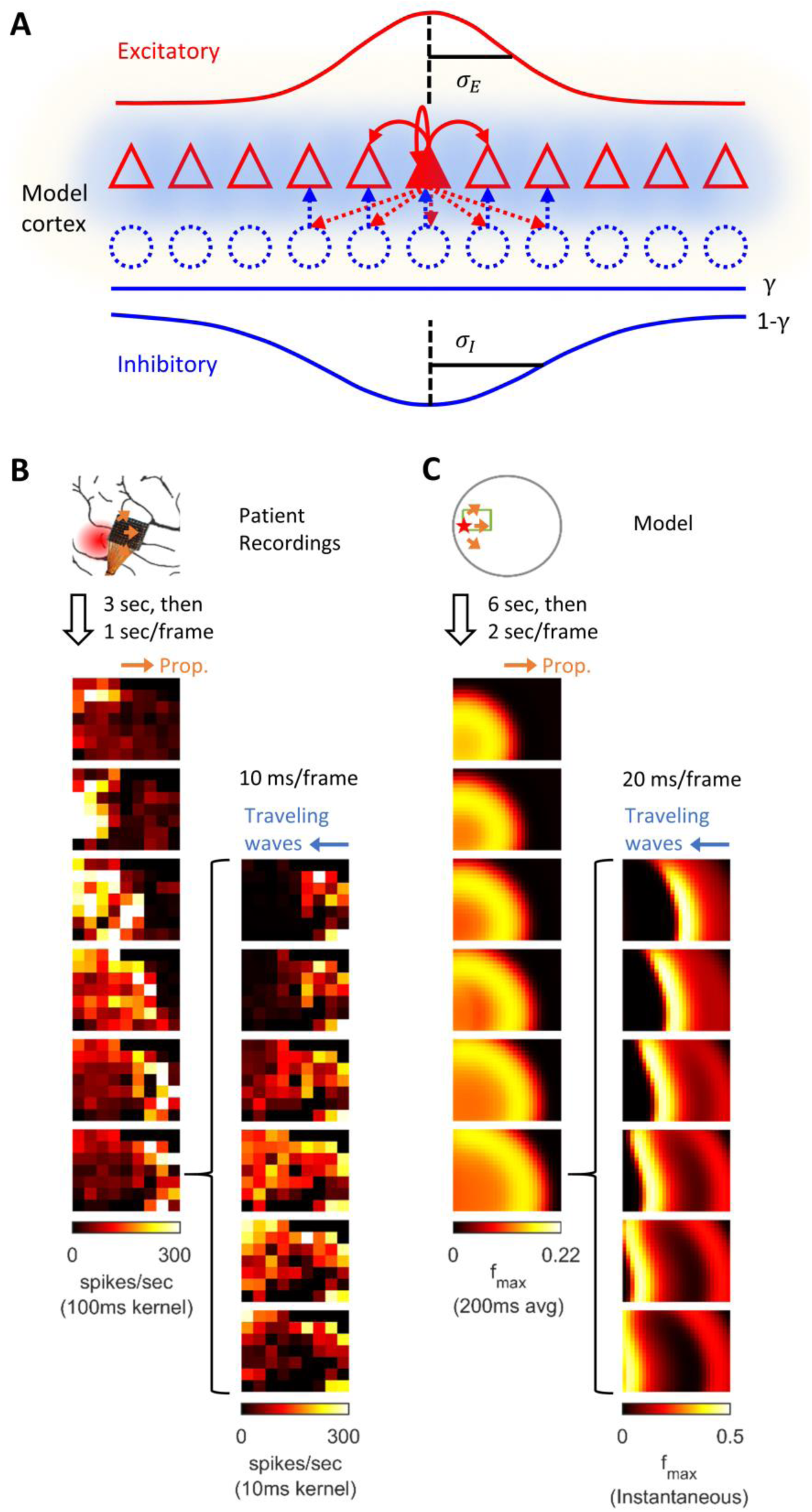
Model schematics and the comparison of patient recordings with model simulation results. A) Model schematic. Triangles: model neurons. Red solid arrows: excitatory recurrents with distance-dependent strength of spatial kernel width σ_*E*_. Dashed blue circles: inhibitory neurons. Dashed blue arrows: distance-dependent recurrent inhibition with spatial kernel width σ_*I*_, which accounts for 1 − *γ* of the total recurrent inhibition. The remaining fraction *γ* of the inhibition is distance independent and represented by the blue hues around model neurons. Interneuron membrane potentials are not explicitly modelled. B) Microelectrode array recordings from Patient A. The red shaded area in the cartoon panel represents the clinically identified seizure onset zone. Orange arrows indicate the seizure propagation direction. The left column shows the spatiotemporal dynamics of multiunit activity in a slow timescale. The right column is a fast timescale zoom-in of the left bottom panel. Evolution of multiunit activity in a slow/fast timescale is estimated by convolving the multiunit spike trains with a 100/10-ms Gaussian kernel respectively. Left-to-right seizure recruitment was seen 3 seconds after the seizure onset (the orange arrow). Once recruited (left bottom panel), fine temporal resolution panels (the right column) showed right-to-left fast waves arose at the edge of the seizure territory (the blue arrow) and traveled toward the internal domain (fast inward traveling waves). C) 2D rate model simulation results. Figure conventions adopted from B. The red star indicates where the seizure-initiating external current input was given (I_d=200pA, duration 3 seconds, covering a round area with radius 5% of the whole neural sheet). Evolution of the neuronal activities within the green rectangle are shown in the panels. Notice the slow outward advancement of seizure territory and the fast-inward traveling waves.

Throughout the manuscript, variations of the primary 2D rate model are introduced to provide more in-depth assessment of key features. Simulation results of the primary model in a noise free environment are first presented and qualitatively compared to patient recordings (Figure 1). Next, the model is simplified to 1-dimensional space to better illustrate physiological mechanisms that account for the slow advancement of the ictal wavefront and the generation of fast inward traveling waves (Figure 2 & 3). A corresponding 1-dimensional spiking model is subsequently shown in Figure 4 & 5 to validate our rate-based approach and explore the effects of spike-timing dependent plasticity on spontaneous seizure occurrence. Finally, the main model is revisited, this time with addition of background current noise, to explain bi-stable seizure evolution endpoints and the origin of complex spatial configurations of focal seizure activities (Figure 6 & 7).

**Figure 2.**
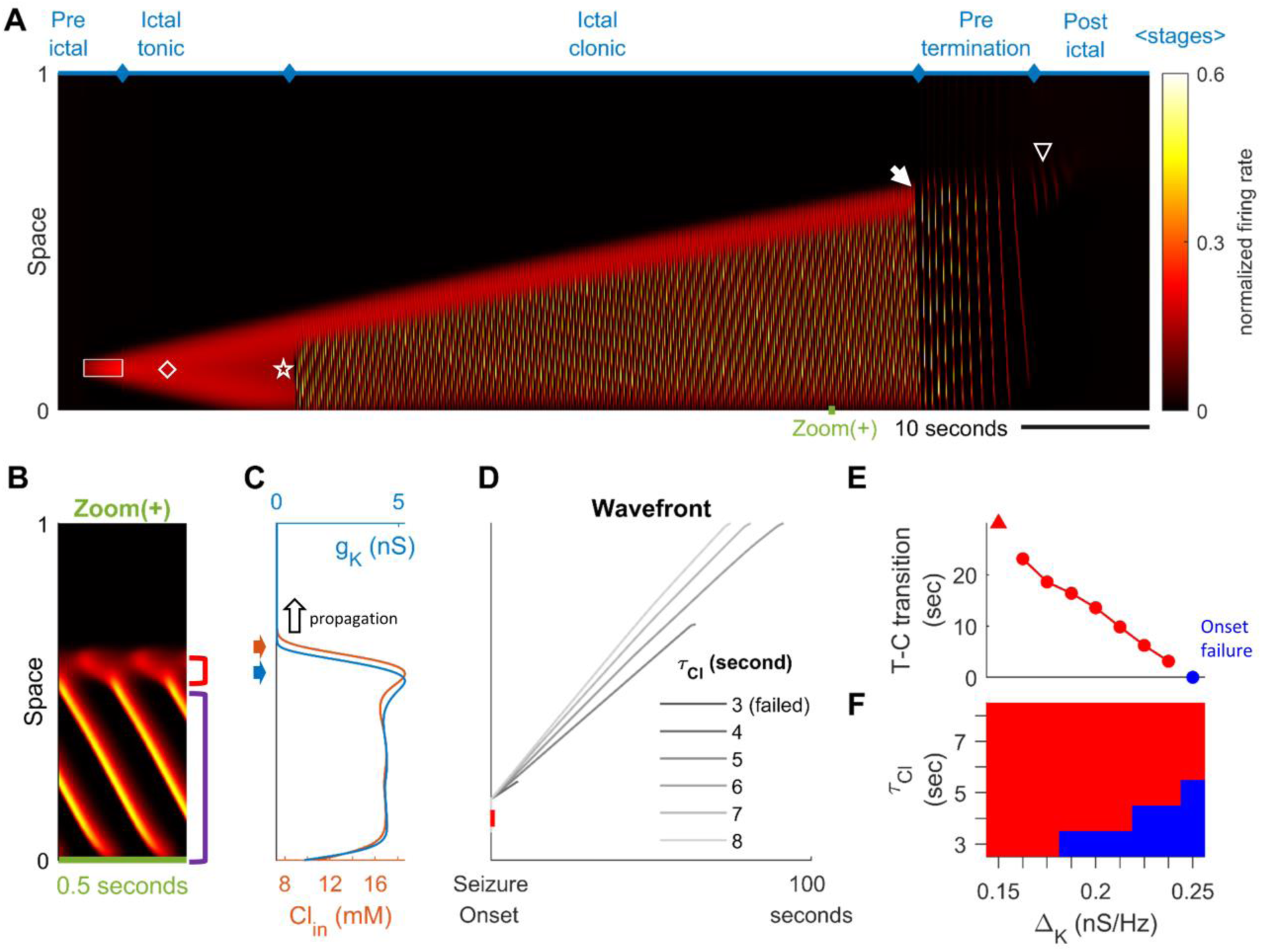
Stages of focal seizure evolution in a 1D model. A) Spatiotemporal evolution of a model seizure in one-dimensional space (*E*_*L*_=-57.5 mV). Horizontal axis: time; vertical-axis: space. Seizure dynamics is partitioned into the following distinct stages: pre-ictal, ictal-tonic, ictal-clonic, pre-termination, and post-ictal. White box: seizure-provoking input (*I*_*d*_=200 pA). White diamond: establishment of external-input-independent tonic-firing area. White star: tonic-to-clonic transition. White arrow: annihilation of the ictal wavefront. White triangle: seizure termination. B) Temporal zoom-in from Panel A during the ictal-clonic stage. Seizure territory can be further subdivided. Red square bracket: the ictal wavefront. Purple square bracket: the internal bursting domain. Note that the fast-moving traveling waves move inwardly. Speed: 1.36 normalized space unit per second. Propagation speed of the ictal wavefront: 0.008 normalized space unit per second, for a traveling wave:wavefront speed ratio of 170. C) Intracellular chloride concentration and sAHP conductance as a function of spatial position during the ictal-clonic stage, corresponding to the activity depicted in panel B. Note the distinct spatial fields of the two processes near the ictal wavefront. D) Impaired chloride clearance allows seizure initiation and speeds up seizure propagation. Gray lines: the expansion of border of the seizure territory versus time. End of the lines: seizure termination, either spontaneous (*τ*_*Cl*_ ≤5 seconds) or after meeting the neural field boundary (*τ*_*Cl*_ ≥6 seconds). E) Activation of the sAHP conductance leads to the tonic-to-clonic transition. Duration of ictal-tonic stage monotonously decreases as the sAHP conductance increases. Red triangle: persistent tonic stage without transition. F) The seizure-permitting area in *τ*_*Cl*_ and Δ_*K*_ parameter space. Blue/red: onset failure/success.

**Figure 3.**
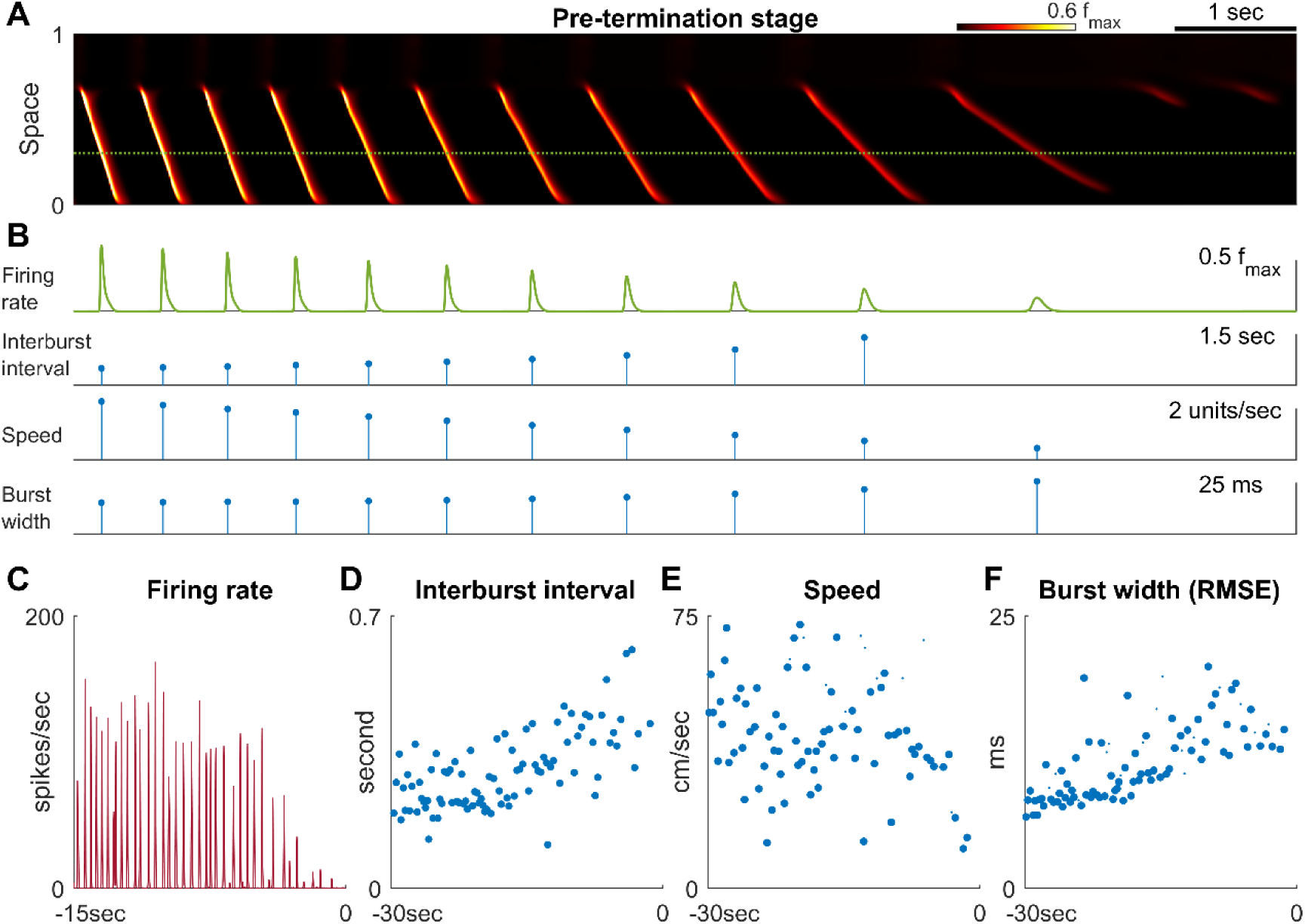
Pre-termination stage shows ‘slowing-down’ dynamics. A) Model seizure, pre-termination stage, zoomed-in from Figure 2A. The horizontal axis is shared with Panel B. The local neuronal population dynamics marked by the dotted green line is further shown in Panel B. Notice that in a space-time diagram a traveling wave is a slant band whose slope is its traveling velocity. B) Quantification of Panel B. Peak firing rate decreases, interburst intervals prolong, traveling wave speed decreases, and burst width decreases as the model seizure approaches termination. Neural activity within a region of 0.05 normalized distance centered at the dotted green line is used to calculate traveling wave speed. Burst width is calculated by first treating *f*(*t*) during each burst (100 ms temporal window) as a distribution and then estimate its standard deviation. C-F) Patient B shows similar trends of neural dynamics at pre-termination stage. Peak firing rate decreased (C) interburst interval increased (D), traveling wave speed decreased (E), and burst width increased (F). Spearman’s correlation coefficients: 0.64 (n=93, p<0.001), −0.62 (p<0.001, n=35), 0.78 (n=93, p<0.001), −0.34 (n=93, p=0.002) for C-F respectively.

**Figure 4.**
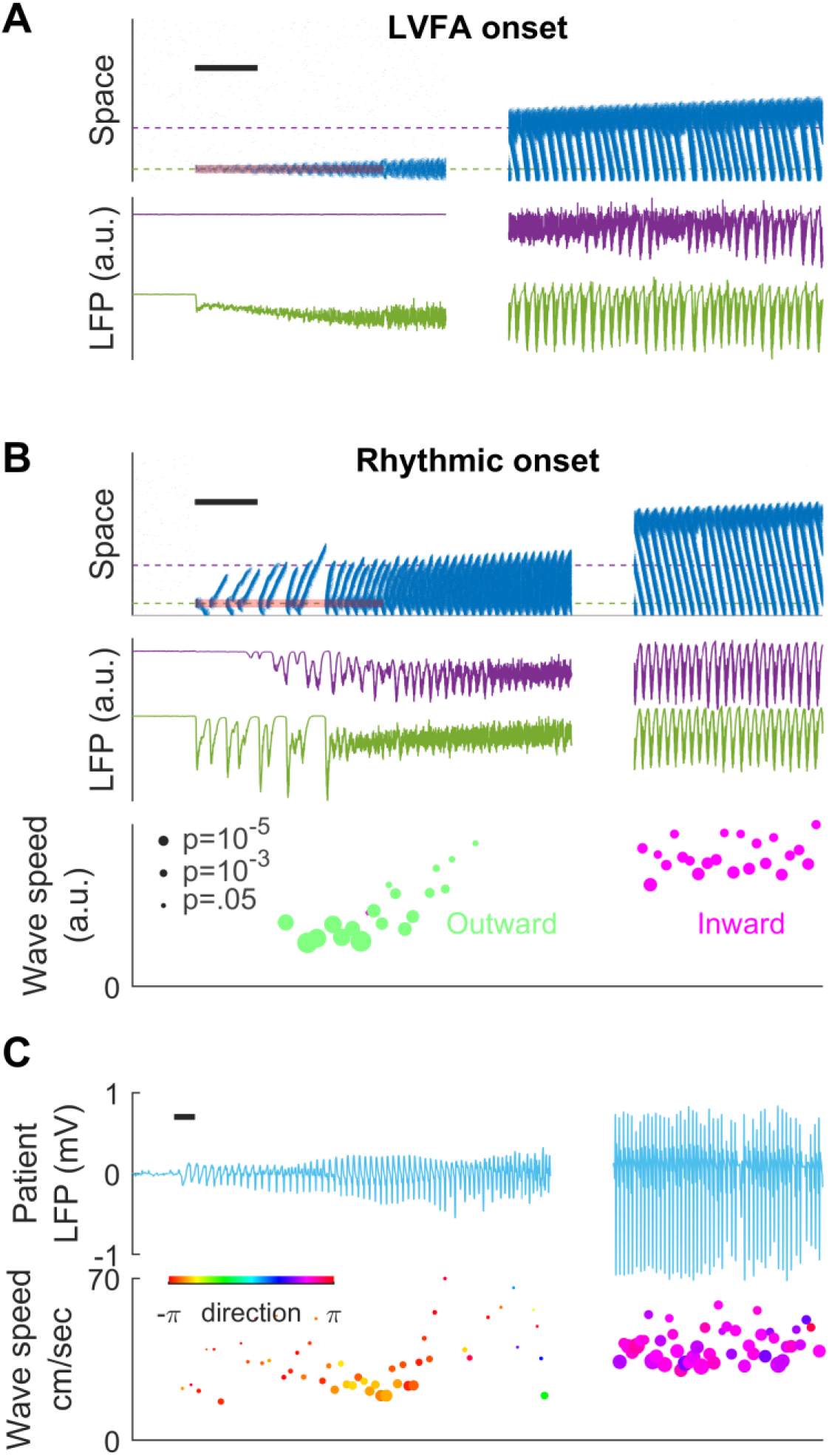
Subtypes of seizure onset pattern can arise from different distributions of recurrent inhibition. A) LVFA onset (*E*_*L*_=-60mV, *γ*=1/6). Black scalebar: 1 second. Upper subpanel: raster plot. Horizontal axis: time. Vertical axis: space. Shaded red zone: *I*_*d*_=80 pA. Green and purple dashed lines indicate where the simulated LFPs, shown in the lower subpanel, are calculated. The blank region is a 20 second interval. Note that LVFA is associated with large DC shift of the LFP, corresponding to seizure onset and ictal wavefront invasion. Periodic LFP discharges emerge after ictal wavefront passage. B) Rhythmic onset (*E*_*L*_=-60mV, *γ*=1/2). Upper and middle subpanels inherit the conventions of panel A. The blank region is a 15 second interval. At seizure onset, several waves are generated and travel outwardly (shaded red area, *I*_*d*_=80pA). They are associated with rhythmic LFP discharges and precede the fast activity. Inward traveling waves emerge after wavefront establishment (right, after the blank interval). Lower subpanel: comparison of speeds between outward and inward traveling waves. Traveling wave speeds are measured locally at the location indicated by the purple dashed line, with each dot’s X-coordinate as the time of local firing rate max. Dot size corresponds to F-test p-value (significance level: 0.001) and color represents the direction of propagation. Outward waves (n=34) travel at a significantly lower speed than inward waves (n=23) (U-test, p<0.001). C) Rhythmic seizure onset recorded from Patient B. Black scalebar: 1 second. Upper subpanel: averaged LFP recorded from the microelectrode array. Lower subpanel: traveling wave direction (dot color) and speed, estimated according to the spatiotemporal distribution of multiunit spikes. The blank region: 40 second interval. The seizure started with periodic LFP discharges before fast activity emerged. As the seizure evolved, traveling wave direction switched (dot color: orange to purple) and the speed increased (U-test p<0.001, n=67 versus 49), analogous to the outward-to-inward switch predicted in Panel B.

**Figure 5.**
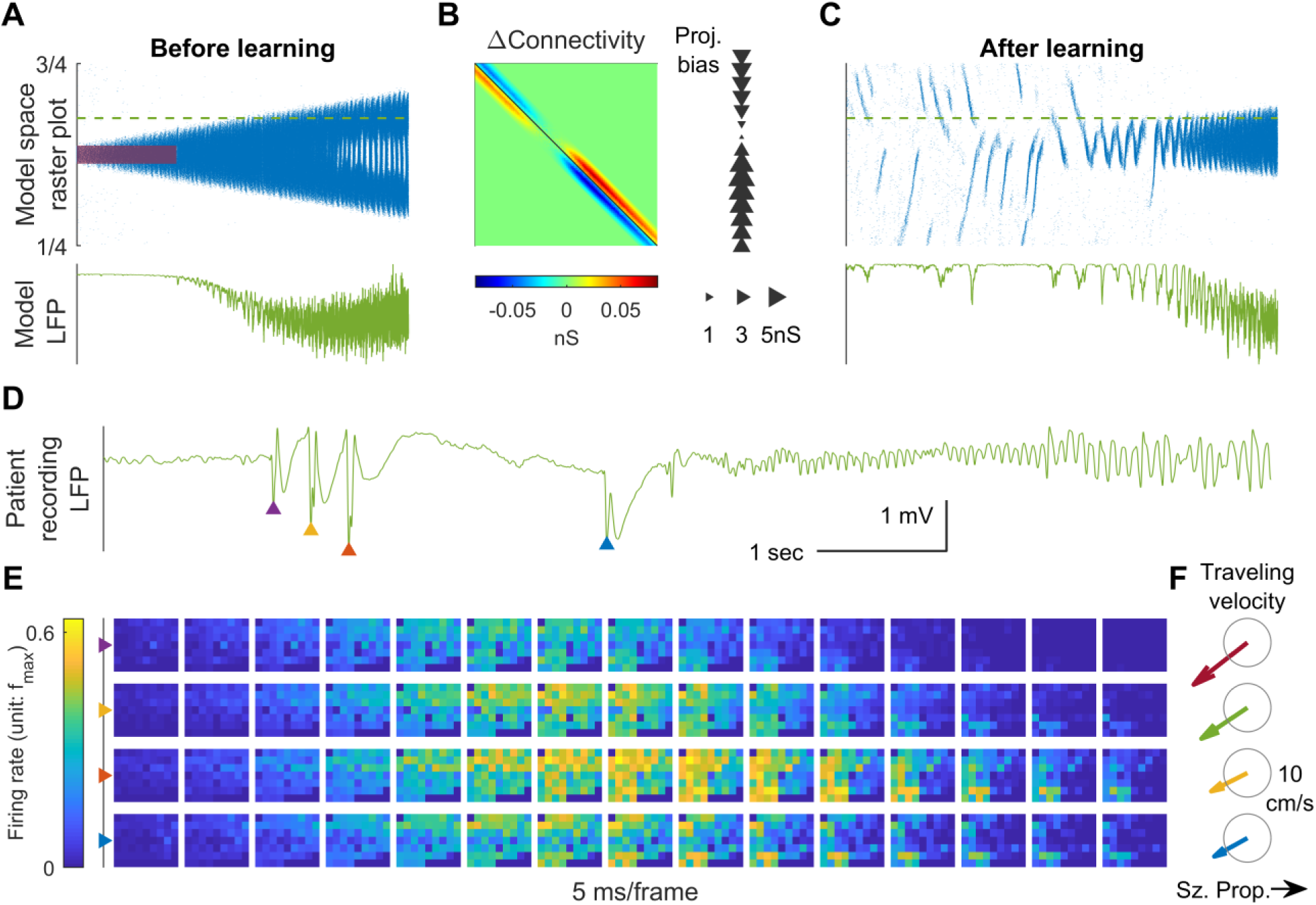
Connectivity induced by spike-timing dependent plasticity may promote emergence of seizure foci. A) Raster plot of a provoked model seizure with LVFA onset pattern. Figure conventions are inherited from Figure 4A. Red shaded area indicates the epileptogenic input (*I*_*d*_=200 pA for 3 seconds). Stochastic background input: σ_*s*_=20 pA, *λ*_*s*_=0.1, *τ*_*s*_=15 ms. The vertical axis is normalized spatial scale and aligned with Panel B & C. The green dashed line indicates where the simulated LFP, shown in the lower subpanel, is calculated. (Δ_*K*_=0.05 nS) B) Changes in recurrent excitatory connectivity strength, *W*_*E*_, induced by STDP after the provoked model seizure (rows correspond to postsynaptic labels, columns to presynaptic). Left subpanel: the Δ*W*_*E*_ matrix. Right subpanel: projection bias (∑_*a*_ Δ*W*_*E,ab*_). The size and direction of the triangles represent the magnitude and direction of the bias respectively. C) A spontaneous seizure develops after STDP learning (*I*_*d*_=0 pA). Several large amplitude LFP discharges, shown in the lower subpanel, precede the seizure onset. These LFP discharges are generated by the centripetal traveling waves seen in the raster plot. D) LFP discharges recorded immediately before Patient A’s seizure onset. Discharges are marked by triangles of different colors. Evolution of the associated multiunit firing pattern is shown in Panel E. E) Multiunit firings constitute traveling waves (10 ms kernel) before LVFA seizure onset. Note that the right half of the array detected multiunit firings earlier than the left half, which is opposite to the expansion direction of the ictal core (left to right). F) Estimated traveling wave direction and speed. Gray circle: 10 cm/sec.

**Figure 6.**
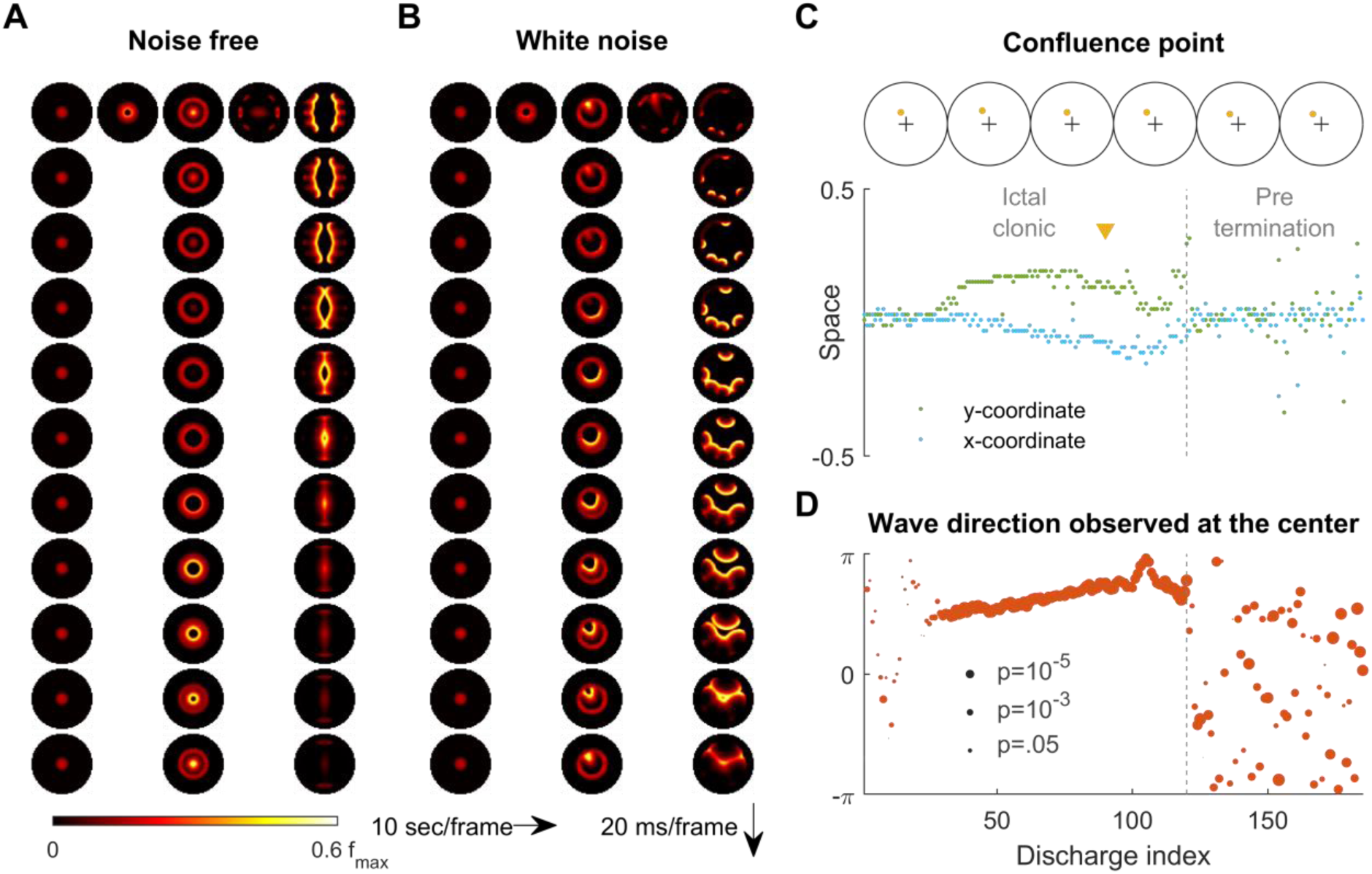
Consistent inward traveling wave direction can emerge from white noise. A) Simulation results of a spatially bounded, two-dimensional rate model under a noise free condition. The epileptogenic stimulus was provided at the center of the round field (*I*_*d*_=200 pA, 3 second, stimulation radius = 0.05). Seizure territory slowly expands (the top row, left to right, 10 seconds per frame) and evolves in stages. The 1^st^ column shows the tonic stage during which the firing rate within the seizure territory stays largely constant (20 ms per frame vertically). The 2^nd^ and 3^rd^ columns show the clonic and pre-termination stages, respectively. Because the neural sheet is symmetric and noise-free, all inward traveling waves meet exactly at the center. B) Simulation protocol as in panel A but with spatiotemporally white noise with diffusion coefficient 200 pA^2^/ms is injected uniformly over the whole neural sheet. The confluence point of inward traveling waves during the clonic stage deviates from the center (middle column). Complex, asymmetric traveling wave generation is observed during the pre-termination stage (right column). C) Locations of confluence points, quantified from Panel B. The upper row: confluence point locations of 6 consecutive inward traveling waves (marked in yellow). The lower subpanel: evolution of the confluence point locations. During the clonic stage (n=120), confluence points significantly deviate from the center (signed-rank test for both *x* and *y* coordinates, p <0.001) and drift slowly (auto-correlation coefficients: *ρ*_*x*_(1)=0.82, *ρ*_*y*_(1)=0.83; both p < 0.001.). D) Traveling wave direction estimated at the center (radius = 0.05). White noise generates a consistent traveling wave direction during the clonic stage once the confluence point drifts away from the center. Circular auto-correlation *ρ*_*θ*_(1) during the ictal clonic stage: 0.97, p<0.001, n=120. However, traveling wave direction becomes more variable during the pre-termination stage: *ρ*_*θ*_(1)=0.09, p=0.5, n=65.

**Figure 7.**
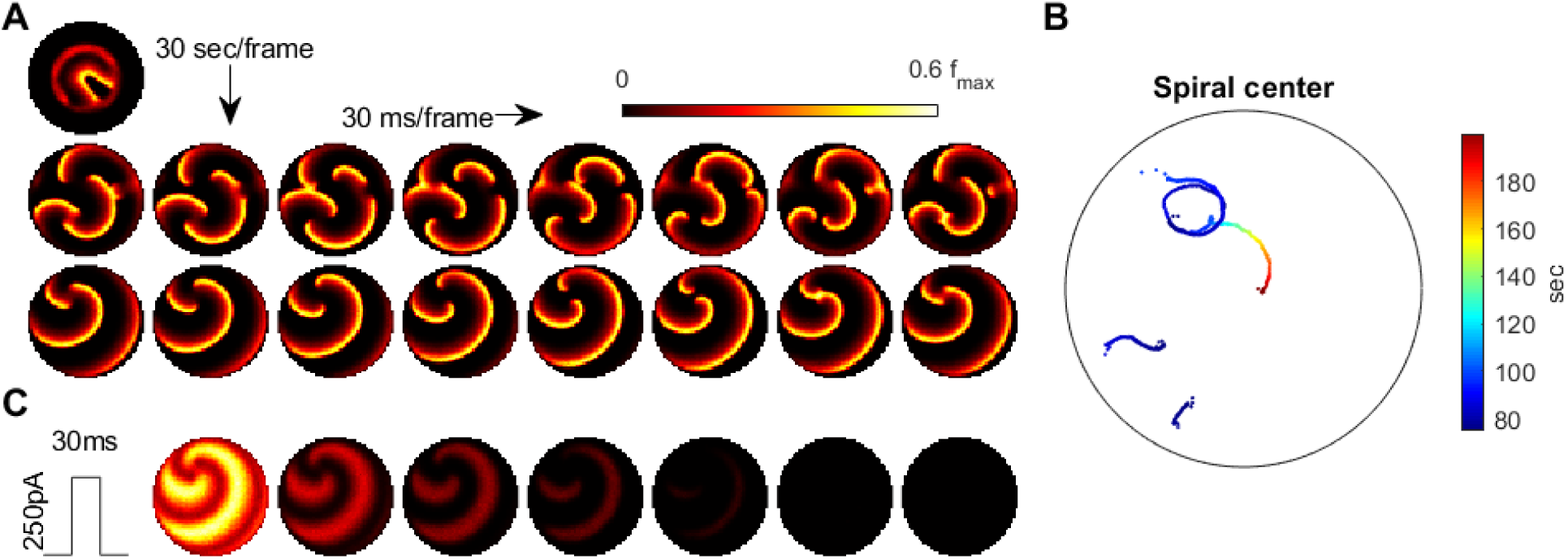
Emergence of spiral waves, status epilepticus, and seizure termination induced by globally synchronizing inputs. A) Spiral wave formation. Model parameters and figure conventions are inherited from Figure 6B. In this simulation, the top single picture shows the ical-clonic stage, then several spiral waves emerge after ictal wavefront annihilation (upper row of pictures). Spiral waves demonstrate complex interactions. Some spiral centers survive and persist indefinitely (lower row). B) Movement of spiral wave centers, quantified from Panel A. One spiral wave eventually dominates the whole field and never terminates. C) A globally synchronizing pulse (*I*_*d*_=250pA, 30 ms) can force spiral-wave seizures to terminate. The pulse is given at the time of the left lower sub-panel in Panel A. Each subpanel is aligned with the lower row of Panel A for comparison.

### Comparison between patient microelectrode recordings and model simulation results

Figure 1B shows a 96-channel Utah array recording from a patient experiencing a typical spontaneous neocortical-onset seizure. The human recording demonstrates a slow, progressive advancement of seizure territory from the left to right side of the array-sampled area (Figure 1B, left, orange arrows). The recruitment was led by the ictal wavefront, which passed through the microelectrode-sampled territory in a few seconds. Following the brief period of tonic firing marking the wavefront location, the neurons, now inside the ictal core, transitioned to repetitive bursting (Figure 1B, right). Bursts propagated sequentially in space from the ictal wavefront back toward the internal domain, constituting fast-moving ‘inward traveling waves’ (Figure 1B, right, the blue arrow). Figure panels 1C shows similar activity patterns generated by our model. In the model, a self-sustaining tonic-firing region was established by a transient focal external stimulus (the red star in Figure 1C). The key dynamics seen in the human recordings were reproduced by the model, including the slow-marching wavefront (Figure 1C, right, orange arrows) and the subsequent fast inward traveling waves (Figure 1C, right, the blue arrow). Supplementary video 1 shows the full spatiotemporal evolution of this model seizure, and unprocessed wideband data of the microelectrode arrays can be found in a previous publication (Schevon *et al*., 2012).

### Stages of focal seizure evolution

To further dissect the main features of seizure dynamics, the 2-dimensional model was reduced into a simplified one-dimensional version (Figure 2). In this model, a transient excitatory input triggers the establishment of a localized, self-sustaining tonic firing region, marking the onset of the model seizure (Figure 2A, white diamond). Within a few seconds, the “activity bump” collapses and transitions spontaneously into repetitive neuronal bursting, marking the transition from the ictal-tonic to the ictal-clonic phase (Figure 2A, white star). The repetitive neuronal bursting happens sequentially according to its distance from the ictal wavefront, forming the fast inward traveling waves (Figure 2B). The fast waves travel two orders of magnitude faster than the speed of the ictal wavefront (Figure 2B). This ratio approximates that previously reported in human recordings, i.e. 0.2 – 0.8 mm/sec for ictal wavefront expansion versus ictal discharge traveling speeds of 20∼50 cm/sec) (Smith *et al*., 2016). As the seizure nears termination, the ictal wavefront dissipates (Figure 2A, white solid arrow), marking the beginning of the pre-termination phase. Bursts become less frequent and begin to spread out and weaken. After the last burst propagates across the territory, the seizure terminates abruptly (Figure 2A, white triangle). These seizure stages and each stage’s key dynamics are further corroborated and validated in the corresponding 1-dimensional integrate-and-fire spiking model (Supplementary Figure S1), that includes threshold noise. The sequence of seizure stages is not affected by duration, size, or intensity of the external seizure-provoking inputs as long as they are adequate to trigger seizure onsets (Supplementary Figure S2).

### Physiology of the ictal wavefront

The slow propagation of the modelled ictal wavefront results from two sequentially activated processes: usage-dependent exhaustion of inhibition and upregulation of adaptation currents (Figure 2C). These are modeled as intracellular chloride accumulation and sAHP conductance activation, respectively.

In our modeled seizures, surround inhibition initially blocks the outward advance of the ictal wavefront. The sequence of events in the transition to seizure is depicted in Figures 2B-C. As the ictal wavefront approaches (Figure 2B, red bracket and Figure 2C, black arrow), intracellular chloride starts to accumulate just ahead of it (brown arrow) due to strong feedforward inhibition projected from the ictal core. Intracellular chloride accumulation causes the cholride reversal potential to become less negative, which compromises the strength of GABA-A receptor-mediated inhibition (Supplementary online methods, equation 3). Inhibition eventually collapses, causing neurons to transition from the resting to the tonic-firing state as they join the ictal wavefront. Intense neuronal firing subsequently activates the sAHP conductance (Figure 2C, blue arrow) (Supplementary online methods, equation 4), which mediates a hyperpolarizing potassium current that curbs neuronal excitability. Consequently, the tonic firing neurons transition into repetitive bursts alternating with periods of neuronal silence, forming the periodic inward traveling waves described above. The sequence of intracellular chloride accumulation followed by the activation of sAHP thus mediates both the slow propagation of the ictal wavefront and the periodic, inward fast traveling waves that characterize the ictal core.

The robustness of inhibition and the amount of adaptation conductance control the degree to which the model is prone to seizures. Model cortices in which neurons can pump out chloride quickly are resistant to seizure invasion. As shown in Figure 2D, speeding up chloride clearance (low *τ_Cl_*) results in a slower ictal wavefront propagation speed, shorter seizure duration, and hence smaller seizure territory. If chloride can be pumped out fast enough, seizure territory cannot even be established (Figure 2D, *τ_Cl_* ≤3 sec). The adaptation conductance (Δ*_K_*) in turn curbs seizures by preventing the establishment of seizure territory and hastening seizure termination. As shown in Figure 2E, increasing the sAHP conductance results in earlier tonic-clonic transitions, earlier seizure termination, and therefore smaller seizure territory. For high levels of the sAHP conductance, seizure initiation fails (Figure 2E, Δ*_K_* ≥0.25nS). Figure 2F shows the seizure-permitting region in the parameter space of *τ_Cl_* and Δ*_K_*. In general, models with fast chloride clearance and strong adaptation conductance are resistant to seizure-provoking inputs.

### Ictal wavefront annihilation and the pre-termination stage

Previously, based on analysis of human multiscale recordings, our group proposed that dissipation of the ictal wavefront is a key event preceding wide-area, simultaneous seizure termination (Smith *et al*., 2016; Liou *et al*., 2017). This process is also evident in the model seizures. As the seizure territory expands, the ictal wavefront in the model encounters increasingly stronger inhibition ahead of itself. This is because non-localized recurrent inhibition (the global part of equation 5 in Supplementary online methods) is progressively activated as more cortical areas are recruited into the seizing territory. The dynamics thus function as a spatial integrator, with strength of inhibition proportional to the extent of the area that has been invaded. Consequentially, seizure propagation gradually slows as the wavefront expands. Once the propagation speed is inadequate to escape from sAHP-induced suppression, the wavefront is annihilated, and the seizure transitions into the pre-termination stage (Figure 2A white triangle & Figure 3).

The pre-termination stage of a model seizure (Figure 3A) shows four characteristic trends: decreasing peak firing rate, increasing inter-burst intervals, decreasing inward traveling wave speed, and increasing burst width (Figure 3B). These ‘slowing-down’ trends match clinical observations as well as the patient recordings (Figure 3C-F). Increased duration of the silent period following each burst allows more time for recovery of inhibition strength, as more chloride can be removed from the intracellular space to restore the transmembrane chloride gradient. Recovered inhibition then attenuates burst intensity (Figure 3C), slows the speed of the inward traveling waves (Figure 3E), and desynchronizes the neuronal population (Figure 3F). The pre-termination stage is characterized by gradually increasing inter-burst intervals (Figure 3A), matching empirical clinical observations as well as patient recordings (Figure 3B). Increased duration of the silent period following each burst allows recovery of inhibition strength, as more chloride can be removed from the intracellular space to restore the transmembrane chloride gradient. Recovered inhibition then attenuates burst intensity (Figure 3A & 3C), desynchronizes the neuronal populations (Figure 3D), and slows the speed of the inward traveling waves (Figure 3E). For the raw electrophysiology signals of traveling waves and methods of determining their velocities, see our previous publications (Smith *et al*., 2016; Liou *et al*., 2017).

### Generalized model for exhaustible inhibition

Although pathological intracellular chloride accumulation is an appealing candidate mechanism for usage-dependent exhaustion of inhibition, other mechanisms have been proposed such as depolarization block of inhibitory neurons (Ahmed *et al*., 2014; Meijer *et al*., 2015), and other pathways that may not be limited to cortical neuronal structures may also exist. We hypothesized that any mechanism that compromises inhibition strength, operates over a time scale of seconds and is triggered by intense inhibition usage is a possible candidate mechanism for the slow expansion of seizure territory and the pre-termination dynamics. To confirm our hypothesis, we modeled a generalized inhibition exhaustion process instead of using the chloride accumulation assumption (Supplementary online methods, equation 8). All evolutionary stages of focal seizures and their characteristic dynamics were reproduced using this generic process (Supplementary Figure S3).

### Distribution of recurrent inhibitory projections determines seizure onset pattern

We have thus far shown that minimal neurophysiologically-based assumptions can account for the complex spatiotemporal evolution of focal seizures. Next, we replaced rate-based units in our one-dimensional model with integrate-and-fire neurons (Supplementary Online Methods, Table 2). LFP-like signals were read out using a computationally validated approximation based on empirical cortical circuit configurations and somatodendritic orientations (Mazzoni et al., 2015). Briefly, local synaptic currents are summed, with the contributions from inhibitory synaptic currents scaled up and delayed in time (Supplementary Online Methods). This allowed us to compare the model LFP with clinical recordings.

Results from the spiking model provide insights into the mechanisms of different focal seizure onset patterns. Low-voltage fast activity (LVFA) is the most common focal seizure onset pattern, with rhythmic discharges or slow rhythmic oscillations seen less frequently (Perucca *et al*., 2014). In our model, seizure onset patterns are determined by the relative strength between local and global recurrent projections (Figure 4 & Supplementary Figure S4). When recurrent projections favor strong localized inhibition, seizure onset is characterized by a localized tonic-firing area, resulting in focal LVFA (Figure 4A, 5:1 ratio of local versus global recurrent inhibition, Supplementary Online Methods, equation 5). The onset pattern shifts to rhythmic discharges when the distribution of recurrent inhibition favors spatially non-specific projections (Figure 4B, 1:1 ratio of local versus global recurrent inhibition, Supplementary Online Methods, equation 5). Under this latter condition, inadequate local restraint allows ictogenic perturbation to spread quickly outward in the form of traveling waves, generating rhythmic discharges as observed in the LFP bands.

Our model predicts that various LFP patterns may coexist during a single seizure episode, depending on regional variances in the distribution of recurrent connectivity (Supplementary Figure S4). Also, our model predicts that outward traveling waves move at a lower speed than inward traveling waves because inhibition outside the ictal wavefront is initially intact (Figure 4B). In agreement with our prediction, as shown in Figure 4C, a similar dependence of traveling wave speed on the direction relative to the ictal wavefront has been observed in human recordings (Smith *et al*., 2016; Liou *et al*., 2017).

### Seizure-induced network remodeling and spontaneous seizure generation

Seizures have been shown to remodel neural networks (Scharfman, 2002; Elger *et al*., 2004; Lenck-Santini and Scott, 2015). We therefore incorporated synaptic plasticity into our model to study how seizures affect network connectivity and, subsequently, seizure dynamics. We subjected the excitatory synaptic projections in our 1D spiking model to spike-timing dependent plasticity (STDP, Supplementary Online method equation 6-7). During a provoked LVFA seizure (Figure 5A), inward traveling waves, because of pre-before-post potentiation in STDP, selectively enhance excitatory projections from the periphery to the internal domain of the ictal core, in accordance with the predominant traveling wave direction (Figure 5B, left). The newly created spatial bias of excitatory projections therefore follows a centripetal pattern (Figure 5B, right, see Supplementary Online method, section Spike-Timing Dependent Plasticity for quantification of spatial biases of synaptic projections). The centripetal connectivity can predispose the neural network to more seizures by funneling background neural activity into the its center. As shown in Figure 5C, a background input which is non-ictogenic in a naïve network may now provoke seizures in the remodeled network by evoking centripetal traveling waves, which can exhaust inhibition strength at the centripetal connectivity center. Subsequent LVFA seizures are therefore prone to be triggered from the same location, and their associated inward traveling waves can further reciprocally strengthen this centripetal connectivity.

Results from seizures generated by our model after the network remodeling described above predict that epileptiform discharges preceding seizure onset travel toward the seizure center, rather than away from it (Figure 5C). Such centripetal traveling waves have been observed in our patient recordings immediately before the onset of an LVFA seizure (Figure 5D-F, compared to Figure 1B). In particular, large-amplitude epileptiform discharges with associated multiunit bursts were present just prior to seizure onset, spreading from right to left (Figures 5E-F). This is opposite to the left-to-right direction of seizure propagation (Figure 1B).

### Variability of traveling wave direction under noisy conditions

We next return to the primary 2D rate model to examine the effects of background noise. Under the noise-free condition considered up to this point, a seizure evoked at the center of the neural sheet expands with a perfectly circular ictal wavefront (Figure 6A, Supplementary Video 2). All inward traveling waves in this case are directed in a centripetal pattern uniformly towards the center point, with no directional preference within the ictal core (Figures 6A, Supplementary Video 2). However, ictal EEG discharges recorded from epilepsy patients clearly demonstrate stable preferred traveling wave directions (Smith *et al*., 2016; Liou *et al*., 2017; Martinet *et al*., 2017). Although preferred directions that correspond to existing functional pathways have been demonstrated (Rossi *et al*., 2017), we found that they may also occur due to random fluctuations in background neural activity.

It might seem that adding random perturbations to our model would make inward traveling waves flip randomly, which would further contradict in vivo observations of a consistent preferred direction in the ictal-clonic stage (Supplementary Figure S5) (Smith *et al*., 2016; Liou *et al*., 2017; Martinet *et al*., 2017). Instead, our model shows that uniformly distributed, spatiotemporally white noise can create a long-lasting, preferred inward traveling wave direction without any requirement for spatial asymmetry or inhomogeneity of the neural sheet (Figure 6B, also see Supplementary Video 3). After the tonic-to-clonic transition in the model, background noise first randomly creates a bias of the confluence point where the inward traveling waves meet (Figure 6C). Once established, the bias persists and, due to the low pass property of the neural sheet, it can only slowly drift, resulting in a sustained traveling wave direction preference lasting until the pre-termination stage. After wavefront annihilation, traveling wave directions randomly fluctuate and flip (Figure 6D). Such variable wave directions mimic human seizure recordings, where an increase in the variability of wave directions and “flip-flop” direction reversals are present before seizure termination (Supplementary Figure S5) (Trevelyan *et al*., 2007a).

### Spiral wave formation, status epilepticus, and synchronization-induced termination

The complex topology of inward traveling waves after ictal wavefront annihilation suggests a candidate scenario that can lead to persistent seizure activity (Figure 7A). We repeated the simulation shown in Figure 6B under the same level of background current noise. Three out of ten repeated simulations showed failure of spontaneous seizure termination. The persistent seizure scenario is characterized by the formation of spiral waves that emerge after wavefront annihilation (Figure 7A) and exhibit complex interactions (Figure 7B). In the example shown, one spiral wave eventually dominated the space, self-attracted and persisted indefinitely, resulting in model status epilepticus (Figure 7B, Supplemental Movie 4). The coexistence of endpoints corresponding to spontaneous seizure or status epilepticus termination under background current noise suggests and may explain the bistable dynamics of seizure termination (Kramer *et al*., 2012).

Sustained spiral-wave activity, i.e. status epilepticus in our model can be terminated by a globally projecting excitatory input that briefly activates the whole neural sheet at once (Figure 7C, Supplementary Movie 4). Within hundreds of milliseconds, the spiral waves spontaneously terminate without going through pre-termination slowing of the discharge pace.

## Discussion

We have presented a biophysically constrained computational model of seizures that, despite being limited to a reduced set of localized dynamic neural properties, reproduces several key aspects of spontaneous human focal seizures that generally manifest on a large scale. We showed that the collapse of inhibition strength ahead of the ictal wavefront, followed by the emergence of hyperpolarizing currents, accounts for the slow expansion of seizure territory, the tonic-to-clonic transition, and the experimentally observed generation of inward traveling waves. These features are closely aligned with properties of human seizures that were previously described using combined clinical EEG and microelectrode recordings (Smith *et al*., 2016). The interplay of usage-dependent inhibition exhaustion and adaptation, which drives wavefront propagation during early stages of the seizure life cycle, also characterizes seizure-prone versus resistant tissues. Our study supports the idea that global recurrent inhibition terminates seizures, creating the slowing pace and traveling wave variability during the pre-termination stage. Additionally, the model demonstrates that distinct seizure onset patterns can result from topologically variant recurrent inhibitory projections. The inclusion of STDP in the model results in progressive, localized pathological enhancement of excitatory connectivity, reducing seizure threshold in the model and enabling spontaneous seizure generation. Our results on plasticity emphasize the importance of the discovery and modeling of rapid inwardly directed traveling waves (Smith *et al*., 2016; Liou *et al*., 2017), as these may induce plasticity that increases susceptibility to future seizures. The model also provides candidate scenarios for both spontaneous seizure termination and ongoing status epilepticus, based on stochastic events following dissipation of the ictal wavefront.

The spatial seizure topology inferred from human multiscale recordings (Schevon *et al*., 2012; Smith *et al*., 2016) and explicitly reproduced in the computational model described here has not previously been studied in its entirety. The propagation of the ictal wavefront is often invisible to standard EEG, and even to wideband microelectrode recordings (Schevon *et al*., 2012). In contrast, the fast-moving inward traveling waves manifest as field potential discharges, forming the classic EEG signature of seizures (Smith *et al*., 2016; Martinet *et al*., 2017). Due to the speed at which these discharges travel (up to 100 cm/sec) (Liou *et al*., 2017), they appear to be synchronized across large brain areas, leading to suppositions of large-area network origin (Kramer *et al*., 2010; Jirsa *et al*., 2014; Khambhati *et al*., 2016). Our model demonstrates that this same appearance can result from well-established neuronal processes occurring at the level of localized cellular interactions. Further, stereotypical traveling wave pathways can result from random fluctuations in background neural activity. A caution applies, however, in interpreting effects that occur near the boundary of the neural sheet, as this area may be artificially hyperexcitable in the model due to reduction in the maximal inhibitory conductances.

Both modern in vitro brain slice studies (Trevelyan *et al*., 2007b) and in vivo human recordings (Schevon *et al*., 2012) support the classical hypothesis of surround inhibition (Prince and Wilder, 1967), which motivated the Mexican-hat pattern of connectivity and the usage-dependent exhaustible inhibition mechanism used in our study (Eissa *et al*., 2017; Liou *et al*., 2018). Theoretically, massive GABAergic activity provoked by ictal events can overwhelm chloride buffering and clearance mechanisms. The collapsed transmembrane chloride gradient compromises the strength of surround inhibition, leading to seizure onset and propagation (Lillis *et al*., 2012; Alfonsa *et al*., 2015). Indeed, excitatory effects of GABAergic transmission have been found in brain slices taken from epilepsy patients (Cohen *et al*., 2002). The paradoxical excitatory effects of GABA in epilepsy patients may be attributed to clearance defects, as tissues from temporal lobe epilepsy (Huberfeld *et al*., 2007) and tumor-associated epilepsy patients (Pallud *et al*., 2014) both showed reduced KCC2 for chloride extrusion. Experimentally knocking down KCC2 also leads to epileptiform discharges (Zhu *et al*., 2005), and an in-vitro fluorescence study showed that chloride accumulation preceded seizure onset (Lillis *et al*., 2012). In agreement with previous studies, our model suggests that tissues with slow chloride clearance are seizure-prone. Inhibition robustness is critical for preventing, restraining, and shaping focal seizure dynamics.

The tonic-clonic transition has previously been theorized to arise from intrinsic neuronal processes such as sAHP (Beverlin *et al*., 2012). In our model, we further explore the hyperpolarizing current’s role throughout a seizure’s life cycle, including controlling seizure onset, mediating tonic-clonic transition, and participating evolution of pre-termination seizure activities. Clinical studies have confirmed the importance of sAHP in prohibiting epileptic activity. Mutations of KCNQ2 and KCNQ3 channels, which contribute to sAHP currents, are associated with familial neonatal epilepsy (Tzingounis and Nicoll, 2008).

In our model, spontaneous seizure termination is caused by progressive build-up of inhibition, which is used to model widespread effect of focal seizures (Burman and Parrish, 2018). This widespread inhibition has not been extensively described, but nevertheless has been demonstrated in both an acute animal seizure model (Liou *et al*., 2018) and in a computational model validated with human microelectrode recordings (Eissa et al., 2017). Experimentally, calcium imaging has confirmed seizure-induced cross-areal activation of PV(+) interneurons, e.g. in visual cortex in response to a focal seizure triggered in somatosensory cortex (Wenzel *et al*., 2017; Liou *et al*., 2018). Anatomically, the effect may be contributed by long-distance, cross-areal projections that preferentially project to inhibitory interneurons in their target circuits (Zhang *et al*., 2014; Sun *et al*., 2019). In addition, focal seizures may depress brain-wide activities via disrupting subcortical structures (Feng et al., 2017) and ascending activation systems (Blumenfeld, 2012). Large-scale non-neural mechanisms that are not specifically modelled in our study may also contribute to seizure termination. For example, global hypoxia secondary to seizure-induced cardiopulmonary compromise may activate ATP-sensitive potassium channels in pyramidal neurons in a spatially non-specific way (Ching et al., 2012), thereby terminating seizures by building resistance ahead of the ictal wavefronts and generate termination dynamics, analogous to the global recurrent inhibition effect modelled in our study. In other words, the global inhibition assumption may be interpreted as a hypothesis that, as a seizure territory enlarges, principle neurons tend to hyperpolarize and are therefore progressively harder to recruit, with the specific neurological mechanisms varying between patients.

In this study, we simplified inhibitory interneurons to reduce model parameter space. This should not be misinterpreted as implying that we consider interneuron dynamics have no role in seizure evolution. Recent studies have shown that interneurons may participate in seizure initiation (Librizzi *et al*., 2017), and their depolarization block can be critical for seizure propagation (Eissa *et al*., 2017). However, complex inhibitory interneuron dynamics, as shown in our study, may not be required for the replication of key features of seizure dynamics.

The biophysical processes proposed here are certainly not the only mechanisms causing seizure evolution. Instead, they should be considered as representative candidates. Processes that produce similar effects on network dynamics and that operate over the same time scales may also contribute to the generation of seizure dynamics, as exemplified by our phenomenological model. For example, in addition to intracellular chloride accumulation, depolarization block of GABAergic neurons may serve a similar role in contributing to inhibition breakdown (Meijer *et al*., 2015). Similarly, aside from hyperpolarizing currents, intense neuronal bursts can inactivate sodium channels, quickly reducing neuronal excitability in a couple of seconds (Fleidervish *et al*., 1996). Additional depressing mechanisms, including excitatory synaptic depletion (Beverlin *et al*., 2012) and inhibitory cell recovery (Ziburkus *et al*., 2006) may collectively contribute to tonic-to-clonic transitions, slowing, and termination. However, key similarities in seizure dynamics observed across different brain regions and pathological conditions suggest that some mechanisms may be universally present, and these are primary candidates for physiological processes underlying seizure evolution.

Our model demonstrates that the spatial distribution of recurrent inhibition can shape neural network dynamics, explaining the variance of focal seizure onset subtypes. Seizures manifesting with focal low voltage fast activity, a positive predictor of seizure freedom following resection of the focus (Alarcon *et al*., 1995), occurred in the setting of relatively strong local inhibition in our model. In contrast, relatively weak local inhibition allows seizure-related activity to spread quickly in the form of periodic outward traveling waves moving ahead of the ictal wavefront (Figure 4B). These outward traveling waves result in the EEG appearance of rhythmic discharging at seizure onset. Previous studies have proposed that this “hypersynchronous” seizure onset pattern depends on long-range cortical connections (Perucca *et al*., 2014; Weiss *et al*., 2016), or increased surrounding tissue excitability (Wang *et al*., 2017). Our model instead highlights the role of localized inhibition and demonstrates that both types of electrographic onset signatures can arise from focal sources, depending primarily on how the ictal foci are established.

There is a complex interplay of seizure dynamics, network remodeling, and epilepsy focus establishment (Mehta *et al*., 1993). In our model, a physiological plasticity mechanism, STDP, is hijacked by the pathological seizure dynamics. The fast inward traveling waves result in self-enhancing pathological centripetal connectivity, resulting in emergence of pre-ictal large amplitude LFP discharges and centripetal waves. Such single or repetitive sharp waves are often seen just prior to LVFA seizure onsets in clinical recordings (Lee *et al*., 2000), and this was also present in one of our human microelectrode recordings. In this recording, the pre-ictal discharges were associated with neurons activated in a centripetal sequence, consistent with our model predictions. Our model also predicts that, within an epileptic focus, seizure dynamics should be highly stereotypical. This has also been supported by previous observations that focal seizure evolution followed fixed sequences, especially during early stages (Jouny *et al*., 2007). Highly-recurrently connected neural clusters have also been found in focal epilepsy patients in imaging studies (Liao *et al*., 2010), suggesting that pathological synaptic connectivity may make neural tissues susceptible to spontaneous seizures. Whether these clusters are seizure sequelae, causes, or both, as suggested in our model, may require further longitudinal studies.

It has been proposed that seizure evolution has bistable endpoints, and that failure of seizure termination can be a stochastic event (Kramer *et al*., 2012). In the presence of noise, the cortex may become trapped in the spiral wave scenario leading to an indefinitely persistent seizure, i.e. status epilepticus. Animal experiments, both in brain slice (Huang *et al*., 2004) and in vivo (Petsche *et al*., 1974; Huang *et al*., 2010) have shown that spiral waves can develop in acute seizure models. Multielectrode array recordings have also revealed spiral epileptiform discharges during feline picrotoxin-induced seizures (Viventi *et al*., 2011). Such spiral waves have yet to be detected in humans, where they may serve as a marker of risk for status epilepticus.

In our model, spiral waves can be terminated by a brief synchronizing input. In agreement with this prediction, synchronization has been shown to promote seizure termination in status epilepticus patients (Schindler *et al*., 2007). Analogously, spiral waves also develop during ventricular fibrillation, a fatal form of cardiac arrhythmia. Delivering a brief pulse that widely activates cardiomyocytes at once has been the standard therapy to rescue such patients. This suggests that a stimulation strategy involving wide, synchronous excitatory input delivery may be an effective approach for seizures characterized by spiral waves.

## Conclusions

We found that a surprisingly small number of well-understood neuronal processes can explain the complex, large-scale spatiotemporal evolution of focal seizures. Our reductionist approach provides insights into generalizable principles underlying complex seizure dynamics, without the need to replicate the seizing brain neuron by neuron. The model’s seizures are notably consistent with clinical presentations and recordings of human seizure events, including the morphology of the ictal wavefront, distinct stages of pre-recruitment, post-recruitment, and pre-termination, wide-ranging ictal and interictal discharges, development of a fixed, chronic seizure focus, spontaneous seizure termination, and status epilepticus. These parallels provide evidence that the topological pattern of a slowly propagating ictal wavefront with rhythmic inward fast traveling waves should be considered the fundamental topology of neocortical focal seizures. Additional investigation is needed to describe the potential impacts of subcortical involvement, cross-hemisphere interaction, and cross-regional propagation.

## Acknowledgments

This work was supported by the National Institutes of Health through National Institute of Neurological Disorders and Stroke grants R01-NS084142 and R01-NS095368, the Gatsby Charitable Trust, the Simons Foundation, and NSF NeuroNex Award DBI-1707398. We thank Steven Siegelbaum, Kenneth Miller, Sean Escola, Ning Qian, and colleagues at Columbia Neurotheory Center for their useful discussions and suggestions.

## Author Contributions

Conceptualization: J.Y.L., L.F.A., C.A.S., and R.G.E. Methodology: J.Y.L. and L.F.A. Software: J.Y.L., C.A.S, S.L.B. Resources and Data Curation (human data): C.A.S., G.M., R.R.G., R.G.E. Writing – Original Draft: J.Y.L. Writing – Review and Editing: J.Y.L., E.H.S., L.M.B., C.A.S., L.F.A. Supervision: L.F.A. and C.A.S. Funding Acquisition: L.F.A. and C.A.S.

## Data Availability Statement

The human seizure data used in this manuscript is currently restricted from open sharing by university policy, but may be shared with qualified investigators under appropriate institutional protections upon request.

## Code Availability Statement

The MATLAB code for the computational model is available at https://github.com/jyunyouliou/LAS-Model

## Online methods

### Data collection and processing

Electrophysiology data were obtained from patients with pharmacoresistant focal epilepsy undergoing invasive EEG monitoring at Columbia University Medical Center as part of their clinical care, and who were additionally enrolled in a study of microelectrode recordings of seizures ^1–3^. The Columbia University Medical Center Institutional Review Board approved the research, and informed consent was obtained from participants prior to surgery. Only data from patients whose microelectrode array-sampled cortical area were recruited into the ictal cores were included in the current study (2 patients, 4 seizures) ^2, 4^. Data from patients who developed generalized seizures or whose array-sampled cortical area lack intense, phase-locked multiunit bursts or propagating ictal wavefront (ictal penumbra) was excluded from this study (5 patients). Data processing algorithms have been previously published. Briefly, multiunit spikes were extracted from Utah array recordings by filtering the raw data between 300 and 3000 Hz and detecting threshold crossings (−4 s.d.) ^5^. Multiunit firing rate was calculated by convolving multiunit spike trains with Gaussian kernels (10 ms for fast dynamics and 100 ms for examining expansion of seizure territory). All calculations were performed using in-house software (Matlab, Mathworks, Natick, MA).

### Rate model

A spatially homogeneous one-dimensional neural field is evenly discretized into 500 populations. Within each population, a principle neuron, described by the following conductance model, is used to approximate population dynamics:

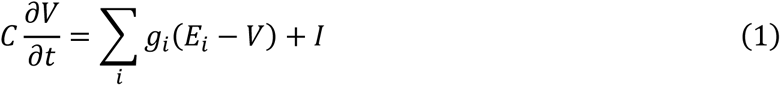

where *V* is membrane potential and *C* is cell capacitance. Four types of conductances (*g*) are modelled: leak (*g_L_*), glutamatergic synaptic (*g_E_*), GABAergic synaptic (*g_I_*), and slow afterhyperpolarization, sAHP, (*g_K_*) conductances. *E_i_* represents each conductance’s reversal potential. Neurons receive external current inputs, *I*, coming from outside the neural network. The inputs are divided into deterministic, *I*_*d*_, and stochastic parts, *I*_*s*_. The stochastic part may be spatiotemporally white or colored, with the latter generated by Ornstein-Uhlenbeck processes with amplitude σ_*s*_ and time and spatial filter constants, *τ*_*s*_ and *λ*_*s*_, respectively. Each population’s mean firing rate, *f*, is calculated by passing the difference between the principle neuron’s membrane potential, *V*, and its firing threshold, *ϕ*, through a sigmoid function, 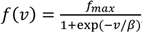, where *υ* = *V* − *ϕ.β* controls slope of the sigmoid function, and *f_max_* is the population’s maximal firing rate. The firing threshold is dynamic with its steady state value linearly dependent on *f* with coefficient Δ_*ϕ*_,

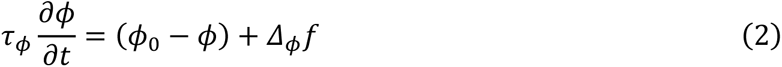

where *ϕ*_0_ represents the principle neuron’s baseline threshold.

The reversal potential of GABAergic conductance (*g*_*I*_) depends on the principle neuron’s transmembrane chloride gradient. Specifically, *E*_*Cl*_ is calculated according to Nernst equation, 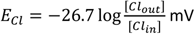. Intracellular chloride concentration is determined by two counteracting mechanisms: chloride current influx and a clearance mechanism with first-order kinetics ^6^,

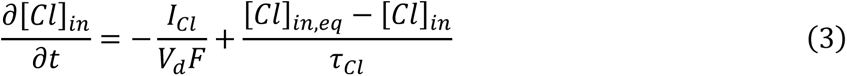

where *I*_*Cl*_ is chloride flow through GABA-A receptors, *I*_*Cl*_ = *g*_*Cl*_(*V* − *E*_*Cl*_), *V*_*d*_ is volume of distribution of intracellular chloride ^7, 8^, *F* is Faraday’s constant, *τ*_*Cl*_ is the time constant of the chloride clearance mechanism ^9^, and [*Cl*]_*in.eq*_ is the equilibrium intracellular chloride concentration. The steady state of the sAHP conductance, *g*_*K*_, is also linearly dependent on firing rate via Δ_*K*_ and evolves according to first-order kinetics,

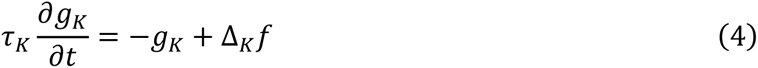

with *τ*_*K*_ is significantly larger than the time constant for threshold adaptation (of order seconds; Table 1).

Inhibitory interneurons are simplified in this study. Their membrane potential dynamics is not specifically modeled, and they react instantly and with a monotonic dependence on their synaptic inputs and only project to their corresponding excitatory neurons. In other words, the model neurons are recurrently connected via “Mexican-hat” connectivity (Figure 1A), emitting both excitatory and inhibitory synaptic projections with the help of these instantly reacting inhibitory interneurons ^10, 11^. Recurrent excitation, *g*_*E*_, is computed by first convolving the normalized firing rate, 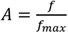, with a spatial kernel, 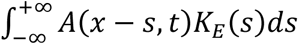, where *K_E_* is a zero-mean Gaussian of variance 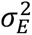. The results are also temporally filtered to model synaptic delay (single exponential with time constant *τ*_*E*_) and then multiplied by the strength of recurrent excitation, 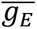. Recurrent inhibition is calculated analogously and can be partitioned into spatially localized and non-localized parts

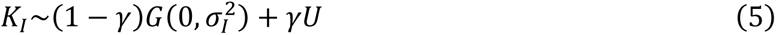

where *G* is a Gaussian, *U* is spatially uniform over the whole neural field and *γ* controls the relative contribution of each component. Recurrent inhibition extends wider than recurrent excitation (σ_*I*_ > σ_*E*_). All models are simulated with *dt*=1 ms. All the model simulation codes are available at https://senselab.med.yale.edu/modeldb/enterCode.cshtml?model=253993.

### Spiking model

2000 neurons with membrane potentials modeled as in equation (1) are evenly distributed along a bounded one-dimensional space. Spikes are emitted stochastically with instantaneous firing rate, 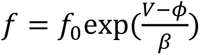, where *f*_0_ is a parameter and *β* quantifies the “uncertainty” of action potential threshold. Numerically, spikes are generated by a Bernoulli process within each time step. If a spike is emitted, the average membrane potential at the spike-emitting time step is taken to be the average of the pre-spiking membrane potential and the peak of the action potential (+40 mV). After a spike, the membrane potential is reset to 20 mV below the pre-spike membrane potential. During the following refractory period (*τ*_*ref*_), membrane potentials are allowed to evolve but spiking is prohibited by setting *f* = 0. In analogy with equation 2, the threshold increases by Δ_*ϕ*_ immediately after a spike, and it exponentially decays with time constant *τ*_*ϕ*_. Chloride dynamics, sAHP conductance, and recurrent projections are modeled as described in the previous section.

### Spike-timing dependent plasticity (STDP)

For simulations of one-dimensional spiking models in which network remodeling is considered, recurrent excitation is subject to spike-timing dependent plasticity (STDP) ^12, 13^. The recurrent synaptic weight matrix, *W*_*E*_, is partitioned into two parts:

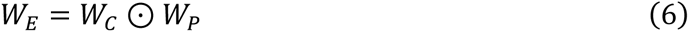

where *W*_*C*_ is the convolution matrix of the Gaussian excitatory kernel (*K*_*E*_), *W*_*P*_ is a matrix representing synaptic weight adjustment according to the STDP rule, and ⊙ represents element-wise multiplication ^14^. Every entry of *W*_*P*_ is initiated at 1. After each model seizure, the plastic part of the synaptic projection from neuron *b* to *a*, namely, the entry *W*_*P,ab*_, is updated according to the STDP rule,

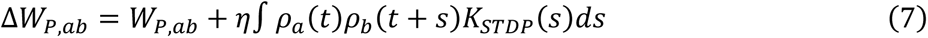

where *ρ* represents the spike trains (delta-functions at the time of the spikes), *η* is the learning rate, and *K*_*STDP*_ is an asymmetric exponential kernel with time constant *τ*_*STDP*_. The learning rate is set so that a single seizure episode maximally changes synaptic weights by 1/3. The STDP-induced connectivity change is reported as Δ*W*_*E*_, and the resultant connectivity is analyzed according to its spatial property. Projections from neuron *b* are divided spatially into two parts: those going to neurons at one side (*W*_*E,ab*_ in which *a* > *b*) and those going to the other side (*a* < *b*). The spatial projection bias of neuron *b* is defined as the difference between the summation of these two parts (∑_*a*<*b*_ *W*_*E,ab*_ − ∑_*a*>*b*_ *W*_*E,ab*_).

### Local field potential (LFP)-like signal readout in the integrate-and-fire spiking model

LFP-like signal is modeled by the reference weighted sum proxy method ^15^. Briefly, LFP is modeled as being proportional to a linear combination of excitatory and synaptic currents, *I*_*E*_ − *αI*_*I*_, where *α* = 1.65 and *I*_*E*_ is temporally delayed by 6 ms, as suggested by Mazzoni et al. Synaptic current contributed by neighboring cells decays exponentially with spatial constant σ_*LFP*_ = 0.025 (unit: normalized spatial scale).

### Generalized model of exhaustible inhibition

The generalized model of exhaustible inhibition is modelled analogously to the rate model except for equation 3. Instead of specifically considering transmembrane chloride dynamics, an abstract dynamic variable, *z*, is used to quantify the effectiveness of inhibition: *g*_*I*_(*x*, *t*) ← *Zg*_*I*_(*x*, *t*). The variable *z* summarizes numerous factors that may contribute to inhibition effectiveness, including chloride gradient, interneuron excitability, short-term plasticity, etc. It is modelled by first-order kinetics

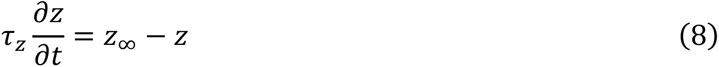

where its steady state, *z*_∞_, depends on the intensity of inhibition usage, *z*_∞_ = *H*(*g*_*I.th*_ − *g*_*I*_), where *H* is a Heaviside step function. When inhibition is strong (above *g*_*I*.*t*ℎ_), its effectiveness decreases exponentially with time constant *τ*_*z*_.

### Two-dimensional model

The 2-D rate model is a based on a bounded, circular-shape two-dimensional space. The simulation neural network partitioned by first generated from a 50 by 50 partitioned square space, and then removing the population that falls outside the round boundary (100 by 100 for high video quality in Supplementary Video 1 and Figure 1). Recurrent connections are calculated by two-dimensional Gaussian convolution in a direct generalization of the 1D case. A round space is used to avoid inhomogeneous boundary effects along specfic directions.

### Quantifying seizure activity in model and patient recordings

For rate models, the duration and spatial involvement of a model seizure is defined as the convex hull constructed from space-time points at which population neuronal firing rate is more than 10% of the maximal firing rate, *Conv* ((***x***, *t*): *f*(***x***, *t*) > 0.1*f*_*max*_). Successful seizure onset is defined as when the provoked seizure activity (*f* > 0.1*f*_*max*_) lasts more than 3 seconds without any requirement of deterministic excitatory current input from outside the neural network. Seizures are partitioned into the following stages: pre-ictal, ictal-tonic (after successful seizure onset), ictal-clonic (after emergence of repetitive bursting activity), pre-termination (after disappearance of all tonic-firing regions), and post-ictal (after all neural population firing rates drop below 0.1 *f_max_*). Transitions between stages are identified visually.

Traveling wave velocities are calculated by least squares linear regression ^16^. For patient data, velocity is inferred by regressing multiunit spike timings against their spatial information (significance level = 0.05) ^16^. For rate models, timings of firing rate peaks are regressed (significance level =10^−3^). Circular standard deviation (20-second window) is used to quantify traveling wave directional variability ^17^.

Spiral waves centers are identified by finding phase singularity points ^18^. At a given time *t*, the instantaneous phase, *ψ*(***x***, *t*), of a population is determined by the state of the membrane potential – threshold plane. *ψ*(***x***, *t*) = *angle*(*V*(***x***, *t*) − *V*_*m*_(***x***) + *i*(*ϕ*(***x***, *t*) − *ϕ*_*m*_(***x***))). *V*_*m*_(***x***) and *ϕ*_*m*_(***x***) are the median voltage and threshold throughout the whole seizure episode at location ***x***. Singular points are found by looking for a closed-path, *l*, that circulates the immediately surrounding points, where ∮_*l*_ ∇*ψ* ⋅ *d****l*** = ±2*π*.

---

**Supplementary Table 1.** Parameters for rate models. Neurons are modelled analogous to neocortical pyramidal neurons in terms of cell capacitance, leak conductance, and membrane time constant ^19^. Maximal synaptic conductances during seizures have been reported in the range of a few hundreds of nS ^20^. Chloride clearance ^9^, buffer ^7^, and seizure-induced sAHP conductance ^21^ are modeled according to the reported ranges. Spatially non-localized recurrent projections (*γ*) are considered in this study as they have been previously reported ^22, 23^. All rate models are based on this standard parameter set. Any further parameter adjustment is reported in figure legends.

| Parameters |  | unit |
| --- | --- | --- |
| $C$ | 100 | pF |
| $g_L$ | 4 | nS |
| $\overline{g_E}$ | 100 | nS |
| $\overline{g_I}$ | 300 | nS |
| $E_L$ | -58 | mV |
| $E_E$ | 0 | mV |
| $E_K$ | -90 | mV |
| $f_{max}$ | 200 | Hz |
| $\beta$ | 2.5 | mV |
| $\tau_E$ | 15 | ms |
| $\tau_I$ | 15 | ms |
| $\tau_\phi$ | 100 | ms |
| $\phi_0$ | -45 | mV |
| $\Delta_\phi$ | 0.3 | mV/Hz |
| $\tau_{Cl}$ | 5 | second |
| $V_d$ | 0.24 | pL |
| $[Cl]_{in.eq}$ | 6 | mM |
| $[Cl]_{out}$ | 110 | mM |
| $\tau_K$ | 5 | second |
| $\Delta_K$ | 0.2 | nS/Hz |
| $\sigma_E$ | 0.02 | Normalized spatial scale |
| $\sigma_I$ | 0.03 | Normalized spatial scale |
| $\gamma$ | 1/6 | |

**Supplementary Table 2.** Parameters for spiking model. Parameters are chosen as described in the legend to Table 1. Time constant of spike timing dependent plasticity is chosen according to previously reported data ^12, 13^.

| Parameters |  | unit |
| --- | --- | --- |
| $C$ | 100 | pF |
| $g_L$ | 4 | nS |
| $g_E$ | 100 | nS |
| $g_I$ | 300 | nS |
| $E_L$ | -57 | mV |
| $E_E$ | 0 | mV |
| $E_K$ | -90 | mV |
| $f_0$ | 2 | Hz |
| $\beta$ | 1.5 | mV |
| $\tau_{ref}$ | 5 | ms |
| $\tau_E$ | 15 | ms |
| $\tau_I$ | 15 | ms |
| $\tau_\phi$ | 100 | mV |
| $\phi_0$ | -55 | mV |
| $\Delta_\phi$ | 2.5 | mV |
| $\tau_{Cl}$ | 5 | Second |
| $V_d$ | 0.24 | pL |
| $[Cl]_{in.eq}$ | 6 | mM |
| $[Cl]_{out}$ | 110 | mM |
| $\tau_K$ | 5 | Second |
| $\Delta_K$ | 0.04 | nS |
| $\sigma_E$ | 0.02 | Normalized spatial scale |
| $\sigma_I$ | 0.03 | Normalized spatial scale |
| $\gamma$ | 1/6 | |
| $\tau_{STDP}$ | 15 | ms |

## References

Ahmed OJ, Kramer MA, Truccolo W, Naftulin JS, Potter NS, Eskandar EN, et al. Inhibitory single neuron control of seizures and epileptic traveling waves in humans. BMC Neuroscience 2014; 15(S1): F3.

Alarcon G, Binnie CD, Elwes RD, Polkey CE. Power spectrum and intracranial EEG patterns at seizure onset in partial epilepsy. Electroencephalogr Clin Neurophysiol 1995; 94(5): 326–37.

Alfonsa H, Merricks EM, Codadu NK, Cunningham MO, Deisseroth K, Racca C, et al. The contribution of raised intraneuronal chloride to epileptic network activity. The Journal of neuroscience: the official journal of the Society for Neuroscience 2015; 35(20): 7715–26.

Beverlin B, 2nd, Kakalios J, Nykamp D, Netoff TI. Dynamical changes in neurons during seizures determine tonic to clonic shift. Journal of computational neuroscience 2012; 33(1): 41–51.

Blumenfeld H. Impaired consciousness in epilepsy. Lancet Neurol 2012; 11(9): 814–26.

Bressloff PC. Waves in Neural Media: From Single Neurons to Neural Fields. Lect N Math Model Li 2014: 1–436.

Buchin A, Chizhov A, Huberfeld G, Miles R, Gutkin BS. Reduced Efficacy of the KCC2 Cotransporter Promotes Epileptic Oscillations in a Subiculum Network Model. The Journal of neuroscience: the official journal of the Society for Neuroscience 2016; 36(46): 11619–33.

Burman RJ, Parrish RR. The Widespread Network Effects of Focal Epilepsy. The Journal of neuroscience: the official journal of the Society for Neuroscience 2018; 38(38): 8107–9.

Ching S, Purdon PL, Vijayan S, Kopell NJ, Brown EN. A neurophysiological-metabolic model for burst suppression. Proceedings of the National Academy of Sciences of the United States of America 2012; 109(8): 3095–100.

Cohen I, Navarro V, Clemenceau S, Baulac M, Miles R. On the origin of interictal activity in human temporal lobe epilepsy in vitro. Science 2002; 298(5597): 1418–21.

Coombes S. Waves, bumps, and patterns in neural field theories. Biological cybernetics 2005; 93(2): 91–108.

Cressman JR, Jr., Ullah G, Ziburkus J, Schiff SJ, Barreto E. The influence of sodium and potassium dynamics on excitability, seizures, and the stability of persistent states: I. Single neuron dynamics. Journal of computational neuroscience 2009; 26(2): 159–70.

Ebersole JS, Husain AM, Nordli DR. Current practice of clinical electroencephalography: Lippincott Williams & Wilkins; 2014.

Eissa TL, Dijkstra K, Brune C, Emerson RG, van Putten M, Goodman RR, et al. Cross-scale effects of neural interactions during human neocortical seizure activity. Proceedings of the National Academy of Sciences of the United States of America 2017; 114(40): 10761–6.

Elger CE, Helmstaedter C, Kurthen M. Chronic epilepsy and cognition. Lancet Neurol 2004; 3(11): 663–72.

Extercatte J, de Haan GJ, Gaitatzis A. Teaching Video NeuroImages: Frontal opercular seizures with jacksonian march. Neurology 2015; 84(11): e83–4.

Feng L, Motelow JE, Ma C, Biche W, McCafferty C, Smith N, et al. Seizures and Sleep in the Thalamus: Focal Limbic Seizures Show Divergent Activity Patterns in Different Thalamic Nuclei. The Journal of neuroscience: the official journal of the Society for Neuroscience 2017; 37(47): 11441–54.

Fleidervish IA, Friedman A, Gutnick MJ. Slow inactivation of Na+ current and slow cumulative spike adaptation in mouse and guinea-pig neocortical neurones in slices. The Journal of physiology 1996; 493 (Pt 1): 83–97.

Huang X, Troy WC, Yang Q, Ma H, Laing CR, Schiff SJ, et al. Spiral waves in disinhibited mammalian neocortex. The Journal of neuroscience: the official journal of the Society for Neuroscience 2004; 24(44): 9897–902.

Huang X, Xu W, Liang J, Takagaki K, Gao X, Wu JY. Spiral wave dynamics in neocortex. Neuron 2010; 68(5): 978–90.

Huberfeld G, Wittner L, Clemenceau S, Baulac M, Kaila K, Miles R, et al. Perturbed chloride homeostasis and GABAergic signaling in human temporal lobe epilepsy. The Journal of neuroscience: the official journal of the Society for Neuroscience 2007; 27(37): 9866–73.

Jirsa VK, Stacey WC, Quilichini PP, Ivanov AI, Bernard C. On the nature of seizure dynamics. Brain: a journal of neurology 2014; 137(Pt 8): 2210–30.

Jouny CC, Adamolekun B, Franaszczuk PJ, Bergey GK. Intrinsic ictal dynamics at the seizure focus: effects of secondary generalization revealed by complexity measures. Epilepsia 2007; 48(2): 297–304.

Khambhati AN, Davis KA, Lucas TH, Litt B, Bassett DS. Virtual Cortical Resection Reveals Push-Pull Network Control Preceding Seizure Evolution. Neuron 2016; 91(5): 1170–82.

Kotagal P, Luders HO. Simple Motor Seizures. In: Engel J, Pedley TA, Aicardi J, editors. Epilepsy: a comprehensive textbook. Second ed: Lippincott Williams & Wilkins; 2008. p. 521–9.

Kramer MA, Eden UT, Kolaczyk ED, Zepeda R, Eskandar EN, Cash SS. Coalescence and fragmentation of cortical networks during focal seizures. The Journal of neuroscience: the official journal of the Society for Neuroscience 2010; 30(30): 10076–85.

Kramer MA, Kirsch HE, Szeri AJ. Pathological pattern formation and cortical propagation of epileptic seizures. J R Soc Interface 2005; 2(2): 113–27.

Kramer MA, Truccolo W, Eden UT, Lepage KQ, Hochberg LR, Eskandar EN, et al. Human seizures self-terminate across spatial scales via a critical transition. Proceedings of the National Academy of Sciences of the United States of America 2012; 109(51): 21116–21.

Lee SA, Spencer DD, Spencer SS. Intracranial EEG seizure-onset patterns in neocortical epilepsy. Epilepsia 2000; 41(3): 297–307.

Lenck-Santini PP, Scott RC. Mechanisms Responsible for Cognitive Impairment in Epilepsy. Cold Spring Harb Perspect Med 2015; 5(10).

Liao W, Zhang Z, Pan Z, Mantini D, Ding J, Duan X, et al. Altered functional connectivity and small-world in mesial temporal lobe epilepsy. PloS one 2010; 5(1): e8525.

Librizzi L, Losi G, Marcon I, Sessolo M, Scalmani P, Carmignoto G, et al. Interneuronal Network Activity at the Onset of Seizure-Like Events in Entorhinal Cortex Slices. The Journal of neuroscience: the official journal of the Society for Neuroscience 2017; 37(43): 10398–407.

Lillis KP, Kramer MA, Mertz J, Staley KJ, White JA. Pyramidal cells accumulate chloride at seizure onset. Neurobiology of disease 2012; 47(3): 358–66.

Liou JY, Ma H, Wenzel M, Zhao M, Baird-Daniel E, Smith EH, et al. Role of inhibitory control in modulating focal seizure spread. Brain: a journal of neurology 2018; 141(7): 2083–97.

Liou JY, Smith EH, Bateman LM, McKhann GM, Goodman RR, Greger B, et al. Multivariate regression methods for estimating velocity of ictal discharges from human microelectrode recordings. Journal of neural engineering 2017; 14(4): 044001.

Martinet LE, Fiddyment G, Madsen JR, Eskandar EN, Truccolo W, Eden UT, et al. Human seizures couple across spatial scales through travelling wave dynamics. Nature communications 2017; 8: 14896.

Mehta MR, Dasgupta C, Ullal GR. A neural network model for kindling of focal epilepsy: basic mechanism. Biological cybernetics 1993; 68(4): 335–40.

Meijer HG, Eissa TL, Kiewiet B, Neuman JF, Schevon CA, Emerson RG, et al. Modeling focal epileptic activity in the Wilson-cowan model with depolarization block. J Math Neurosci 2015; 5: 7.

Pallud J, Le Van Quyen M, Bielle F, Pellegrino C, Varlet P, Labussiere M, et al. Cortical GABAergic excitation contributes to epileptic activities around human glioma. Sci Transl Med 2014; 6(244): 244ra89.

Perucca P, Dubeau F, Gotman J. Intracranial electroencephalographic seizure-onset patterns: effect of underlying pathology. Brain: a journal of neurology 2014; 137(Pt 1): 183–96.

Petsche H, Prohaska O, Rappelsberger P, Vollmer R, Kaiser A. Cortical seizure patterns in multidimensional view: the information content of equipotential maps. Epilepsia 1974; 15(4): 439–63.

Prince DA, Wilder BJ. Control mechanisms in cortical epileptogenic foci. “Surround” inhibition. Arch Neurol 1967; 16(2): 194–202.

Proix T, Jirsa VK, Bartolomei F, Guye M, Truccolo W. Predicting the spatiotemporal diversity of seizure propagation and termination in human focal epilepsy. Nature communications 2018; 9(1): 1088.

Rossi LF, Wykes RC, Kullmann DM, Carandini M. Focal cortical seizures start as standing waves and propagate respecting homotopic connectivity. Nature communications 2017; 8(1): 217.

Scharfman HE. Epilepsy as an example of neural plasticity. The Neuroscientist: a review journal bringing neurobiology, neurology and psychiatry 2002; 8(2): 154–73.

Schevon CA, Weiss SA, McKhann G, Jr., Goodman RR, Yuste R, Emerson RG, et al. Evidence of an inhibitory restraint of seizure activity in humans. Nature communications 2012; 3: 1060.

Schindler K, Elger CE, Lehnertz K. Increasing synchronization may promote seizure termination: evidence from status epilepticus. Clin Neurophysiol 2007; 118(9): 1955–68.

Smith EH, Liou JY, Davis TS, Merricks EM, Kellis SS, Weiss SA, et al. The ictal wavefront is the spatiotemporal source of discharges during spontaneous human seizures. Nature communications 2016; 7: 11098.

Soltesz I, Staley K. Computational neuroscience in epilepsy: Academic Press; 2011.

Sun Q, Li X, Ren M, Zhao M, Zhong Q, Ren Y, et al. A whole-brain map of long-range inputs to GABAergic interneurons in the mouse medial prefrontal cortex. Nature neuroscience 2019; 22(8): 1357–70.

Trevelyan AJ. The direct relationship between inhibitory currents and local field potentials. The Journal of neuroscience: the official journal of the Society for Neuroscience 2009; 29(48): 15299–307.

Trevelyan AJ, Baldeweg T, van Drongelen W, Yuste R, Whittington M. The source of afterdischarge activity in neocortical tonic-clonic epilepsy. The Journal of neuroscience: the official journal of the Society for Neuroscience 2007a; 27(49): 13513–9.

Trevelyan AJ, Sussillo D, Watson BO, Yuste R. Modular propagation of epileptiform activity: evidence for an inhibitory veto in neocortex. The Journal of neuroscience: the official journal of the Society for Neuroscience 2006; 26(48): 12447-55.

Trevelyan AJ, Sussillo D, Yuste R. Feedforward inhibition contributes to the control of epileptiform propagation speed. The Journal of neuroscience: the official journal of the Society for Neuroscience 2007b; 27(13): 3383–7.

Tzingounis AV, Nicoll RA. Contribution of KCNQ2 and KCNQ3 to the medium and slow afterhyperpolarization currents. Proceedings of the National Academy of Sciences of the United States of America 2008; 105(50): 19974–9.

Ullah G, Cressman JR, Jr., Barreto E, Schiff SJ. The influence of sodium and potassium dynamics on excitability, seizures, and the stability of persistent states. II. Network and glial dynamics. Journal of computational neuroscience 2009; 26(2): 171–83.

Viventi J, Kim DH, Vigeland L, Frechette ES, Blanco JA, Kim YS, et al. Flexible, foldable, actively multiplexed, high-density electrode array for mapping brain activity in vivo. Nature neuroscience 2011; 14(12): 1599–605.

Wang Y, Trevelyan AJ, Valentin A, Alarcon G, Taylor PN, Kaiser M. Mechanisms underlying different onset patterns of focal seizures. PLoS computational biology 2017; 13(5): e1005475.

Weiss SA, Alvarado-Rojas C, Bragin A, Behnke E, Fields T, Fried I, et al. Ictal onset patterns of local field potentials, high frequency oscillations, and unit activity in human mesial temporal lobe epilepsy. Epilepsia 2016; 57(1): 111–21.

Wenzel M, Hamm JP, Peterka DS, Yuste R. Reliable and Elastic Propagation of Cortical Seizures In Vivo. Cell Rep 2017; 19(13): 2681–93.

Worthington M. Diagnoses in Assyrian and Babylonian medicine: ancient sources, translations, and modern medical analyses. Med Hist 2007; 51(2): 269–71.

Yang KH, Franaszczuk PJ, Bergey GK. Inhibition modifies the effects of slow calcium-activated potassium channels on epileptiform activity in a neuronal network model. Biological cybernetics 2005; 92(2): 71–81.

York GK, 3rd, Steinberg DA. Hughlings Jackson’s neurological ideas. Brain: a journal of neurology 2011; 134(Pt 10): 3106–13.

Zhang S, Xu M, Kamigaki T, Hoang Do JP, Chang WC, Jenvay S, et al. Selective attention. Long-range and local circuits for top-down modulation of visual cortex processing. Science 2014; 345(6197): 660–5.

Zhu L, Lovinger D, Delpire E. Cortical neurons lacking KCC2 expression show impaired regulation of intracellular chloride. Journal of neurophysiology 2005; 93(3): 1557–68.

Ziburkus J, Cressman JR, Barreto E, Schiff SJ. Interneuron and pyramidal cell interplay during in vitro seizure-like events. Journal of neurophysiology 2006; 95(6): 3948–54.

## References

1. Schevon, C.A., et al. Microphysiology of epileptiform activity in human neocortex. J Clin Neurophysiol 25, 321–330 (2008).

2. Schevon, C.A., et al. Evidence of an inhibitory restraint of seizure activity in humans. Nature communications 3, 1060 (2012).

3. Waziri, A., et al. Initial surgical experience with a dense cortical microarray in epileptic patients undergoing craniotomy for subdural electrode implantation. Neurosurgery 64, 540–545; discussion 545 (2009).

4. Smith, E.H., et al. The ictal wavefront is the spatiotemporal source of discharges during spontaneous human seizures. Nature communications 7, 11098 (2016).

5. Quiroga, R.Q., Nadasdy, Z. & Ben-Shaul, Y. Unsupervised spike detection and sorting with wavelets and superparamagnetic clustering. Neural computation 16, 1661–1687 (2004).

6. Zhu, L., Lovinger, D. & Delpire, E. Cortical neurons lacking KCC2 expression show impaired regulation of intracellular chloride. Journal of neurophysiology 93, 1557–1568 (2005).

7. Marchetti, C., Tabak, J., Chub, N., O’Donovan, M.J. & Rinzel, J. Modeling spontaneous activity in the developing spinal cord using activity-dependent variations of intracellular chloride. The Journal of neuroscience: the official journal of the Society for Neuroscience 25, 3601–3612 (2005).

8. Vladimirski, B.B., Tabak, J., O’Donovan, M.J. & Rinzel, J. Episodic activity in a heterogeneous excitatory network, from spiking neurons to mean field. Journal of computational neuroscience 25, 39–63 (2008).

9. Deisz, R.A., Lehmann, T.N., Horn, P., Dehnicke, C. & Nitsch, R. Components of neuronal chloride transport in rat and human neocortex. The Journal of physiology 589, 1317–1347 (2011).

10. Dayan, P. & Abbott, L.F. Theoretical neuroscience: computational and mathematical modeling of neural systems, (MIT Press, Cambridge, Mass., 2001).

11. Bressloff, P.C. Waves in Neural Media: From Single Neurons to Neural Fields. Lect N Math Model Li, 1–436 (2014).

12. Bi, G.Q. & Poo, M.M. Synaptic modifications in cultured hippocampal neurons: dependence on spike timing, synaptic strength, and postsynaptic cell type. The Journal of neuroscience: the official journal of the Society for Neuroscience 18, 10464–10472 (1998).

13. Song, S., Miller, K.D. & Abbott, L.F. Competitive Hebbian learning through spike-timing-dependent synaptic plasticity. Nature neuroscience 3, 919–926 (2000).

14. Toyoizumi, T., Kaneko, M., Stryker, M.P. & Miller, K.D. Modeling the dynamic interaction of Hebbian and homeostatic plasticity. Neuron 84, 497–510 (2014).

15. Mazzoni, A., et al. Computing the Local Field Potential (LFP) from Integrate-and-Fire Network Models. PLoS computational biology 11, e1004584 (2015).

16. Liou, J.Y., et al. Multivariate regression methods for estimating velocity of ictal discharges from human microelectrode recordings. Journal of neural engineering (2017).

17. Berens, P. CircStat: A MATLAB Toolbox for Circular Statistics. J Stat Softw 31, 1–21 (2009).

18. Iyer, A.N. & Gray, R.A. An experimentalist’s approach to accurate localization of phase singularities during reentry. Ann Biomed Eng 29, 47–59 (2001).

19. Tripathy, S.J., Savitskaya, J., Burton, S.D., Urban, N.N. & Gerkin, R.C. NeuroElectro: a window to the world’s neuron electrophysiology data. Frontiers in neuroinformatics 8, 40 (2014).

20. Neckelmann, D., Amzica, F. & Steriade, M. Changes in neuronal conductance during different components of cortically generated spike-wave seizures. Neuroscience 96, 475–485 (2000).

21. Alger, B.E. & Nicoll, R.A. Epileptiform burst afterhyperolarization: calcium-dependent potassium potential in hippocampal CA1 pyramidal cells. Science 210, 1122–1124 (1980).

22. Braitenberg, V. & Schüz, A. Cortex: statistics and geometry of neuronal connectivity, (Springer Science & Business Media, 2013).

23. Liou, J.Y., et al. Role of inhibitory control in modulating focal seizure spread. Brain: a journal of neurology 141, 2083–2097 (2018).

