## Supplementary Figures for "A theoretical model for focal seizure initiation, propagation, termination, and progression"

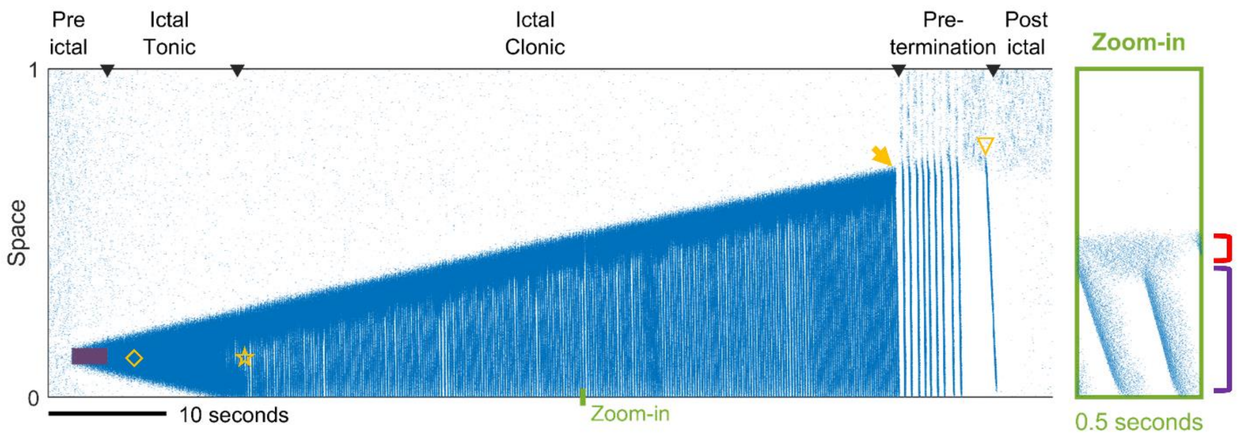


Supplementary Figure S1. Stages of seizure evolution in a spiking model

Left) Spatiotemporal evolution of a model seizure simulated in a one-dimensional spiking neural network. The spiking network is set to be similar to the rate model shown in Figure 2A ($E_L$=-57.25 mV, $\beta$=2.5 mV). Analogous to Figure 2A, seizure dynamics can be partitioned into distinct stages: pre-ictal, ictal-tonic, ictal-clonic, pre-termination, and post-ictal. Red shaded area: deterministic external excitatory input, $I_d$=200 pA. Yellow diamond: establishment of external input-independent tonic-firing area. Yellow star: tonic-to-clonic transition. After tonic-to-clonic transition, seizure territory can be partitioned into the tonic-firing edge (wavefront) and internal bursting domain (inward traveling waves). Yellow arrow: annihilation of the ictal wavefront. Yellow triangle: seizure termination.

Right) Temporal zoom-in from the left panel during the ictal-clonic stage. Red square bracket: ictal wavefront. Purple square bracket: internal bursting domain.

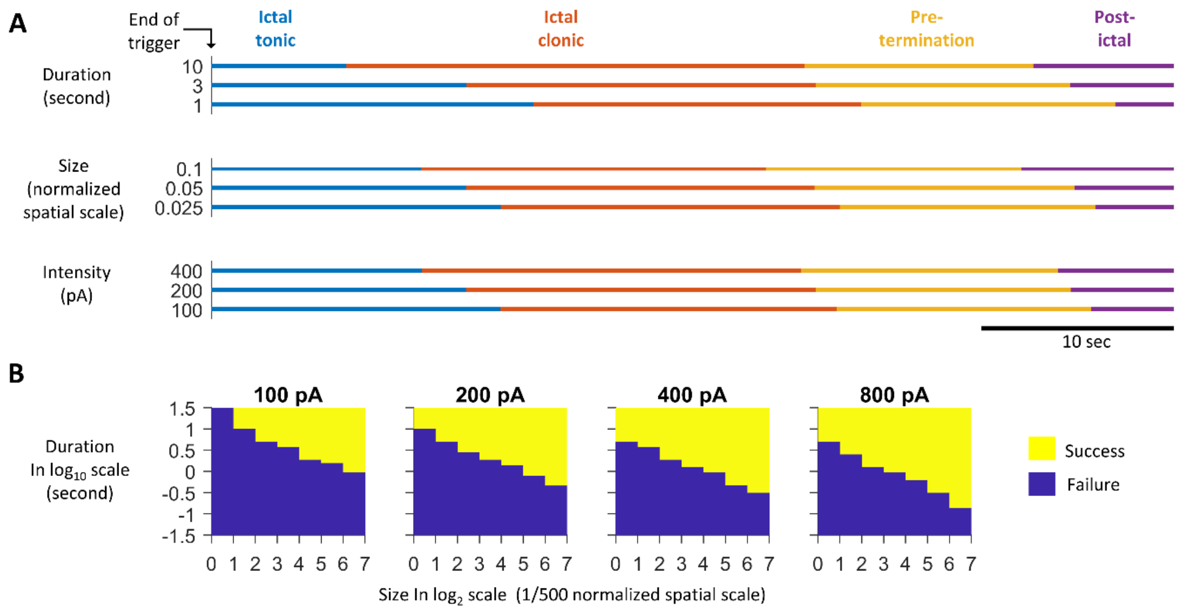


Supplementary Figure S2. Effects of input triggers on model seizure dynamics.

A) Varying duration, size, and intensity of seizure-provoking inputs does not alter the sequence of seizure stages. The external inputs, $I(x,t)$, are provided to the one-dimensional rate model at the center of the space.

B) High input currents, long duration, and large size focal inputs are more effective at triggering seizures. Successful seizure onset is defined if the activity of any rate unit is > 0.1 $f_{max}$ 5 seconds after the end of the external input trigger. Parameter search was done over a logarithm scale grid. Minimal X-axis grid = 1 and Y-axis grid = 1/8.

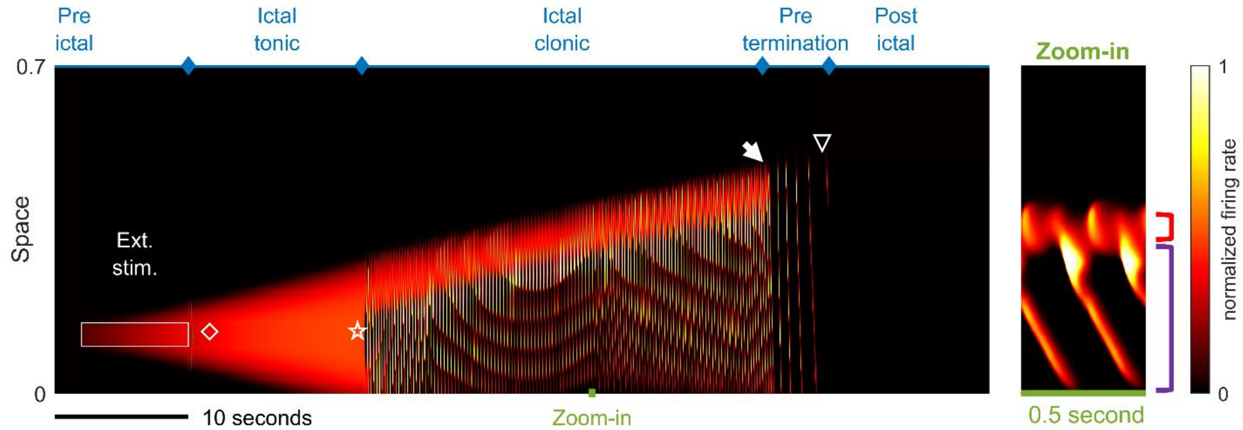


Supplementary Figure S3. Seizure evolution in the generalized model of exhaustible inhibition

Figure conventions inherited from Figure 2A. All stages of seizure evolution and their characteristic dynamics can be reproduced in the generalized model. Parameters are adopted from Table 1 with the following adjustments: $\Delta_K$=0.125 nS/Hz; $\tau_K$=7 seconds; $g_{I.th}$=15 nS; $\overline{g_E}$=125 nS; $E_L$=-60 mV. Deterministic external current input: $I_d$=200 pA.

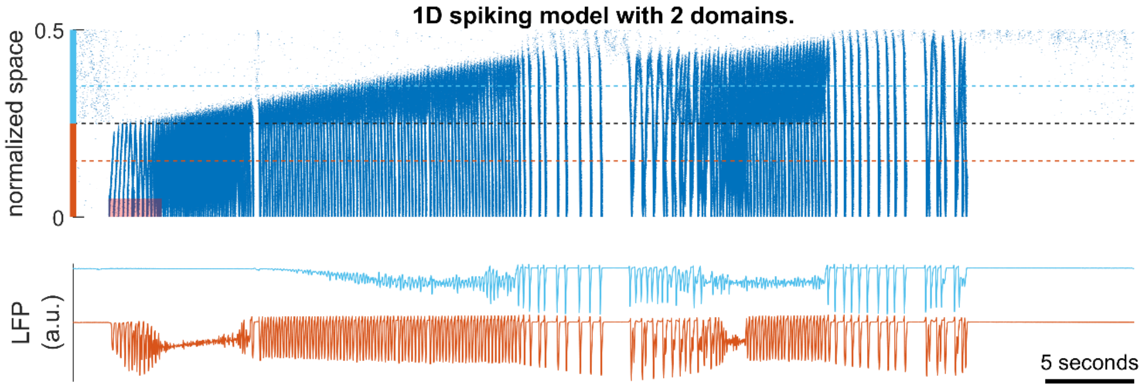


Supplementary Figure S4. Local environment determines seizure onset, propagation and termination patterns

Upper panel: raster plot of a model seizure simulated in a spiking network with two domains (light blue and orange). The upper part of the space (light blue) is configured as Supplementary Figure S1 ($\gamma$=1/2), whereas the lower part (orange) is configured as Figure 4B ($\gamma$=1/6). The gray dashed line indicates the boundary between the two domains. The light blue and orange dashed lines indicate where the simulated LFPs, shown in the lower subpanel, are calculated. The shaded red area: $I_d$=80 pA. Outward traveling waves with rhythmic LFP discharges can be seen at the lower domain; whereas LFP simulated in the upper domain shows LVFA soon after seizure onset. The seizure approaches termination two times but fails the first time. Notice the characteristic slowing pre-termination dynamics is associated with inward traveling waves arising from the upper domain.

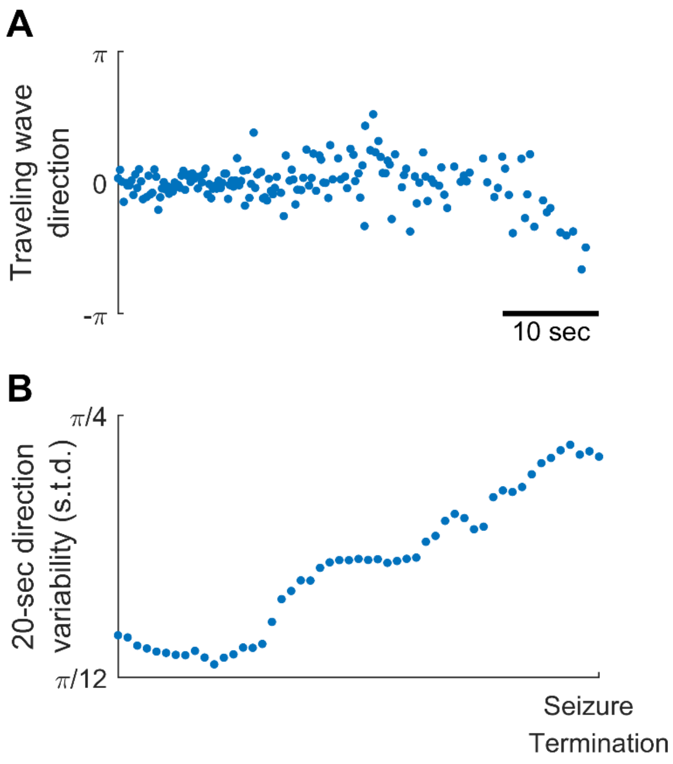


Supplementary Figure S5. Increasing traveling wave direction variability as Patient B's seizure approached termination.

A) Directions of Patient B's ictal traveling waves before seizure termination (the same ones as shown in Figure 3). Horizontal axis: time. The axis is shared with Panel B.
B) Evolution of traveling direction variability estimated by a 20-second moving window, plotted every 1 second, quantified by circular standard deviation. Traveling wave direction variability increases as the seizure approaches termination (Spearman's correlation coefficient: 0.95, $p<0.001$, $n=50$).
