## Supplementary Video Legends for "A theoretical model for focal seizure initiation, propagation, termination, and progression"

Supplementary Video 1.

Full evolution of the model seizure shown in Figure 1C.

Supplementary Video 2.

Full evolution of the model seizure shown in Figure 6A.

Supplementary Video 3.

Full evolution of the model seizure shown in Figure 6B.

Supplementary Video 4

Full evolution of the model seizure shown in Figure 7.
